# Land-use and related pressures have reduced biotic integrity more on islands than on mainlands

**DOI:** 10.1101/576546

**Authors:** Katia Sanchez-Ortiz, Ricardo E. Gonzalez, Adriana De Palma, Tim Newbold, Samantha L. L. Hill, Jason M. Tylianakis, Luca Börger, Igor Lysenko, Andy Purvis

## Abstract

Tracking progress towards biodiversity targets requires indicators that are sensitive to changes at policy-relevant scales, can easily be aggregated to any spatial scale and are simple to understand. The Biodiversity Intactness Index (BII), which estimates the average abundance of a diverse set of organisms in a given area relative to their reference populations, was proposed in 2005 in response to this need. A new implementation of BII was developed as part of the PREDICTS project in 2016 and has been adopted by GEO BON, IPBES and CBD. The previous global models for BII estimation could not account for pressures having different effects in different settings. Islands are a setting of particular interest: many are home to a disproportionate number of endemic species; oceanic islands may have relatively low overall species diversity because of their isolation; and the pattern and timing of human pressures can be very different from that seen on mainlands. Here, we test whether biotic integrity – as estimated by BII – has decreased more severely on islands than mainlands. We update methods previously used to estimate BII globally (Newbold et al., 2016) to allow pressure effects to differ between islands and mainlands, while also implementing some other recent improvements in modelling. We estimate BII for islands and mainlands by combining global models of how two aspects of biodiversity – overall abundance, and compositional similarity to minimally-impacted sites – have been affected by human pressures. We use these models to project high-resolution (∼1km^2^) global maps of BII for the year 2005. We calculate average BII for island and mainland biomes, countries, IPBES regions and biodiversity hotspots; and repeat our analyses using a richness-based version of BII. BII on both islands and mainlands has fallen below the values proposed as safe limits across most regions, biomes and biodiversity hotspots. Our BII estimates are lower than those published in 2016, globally, within all biodiversity hotspots and within most biomes, and show greater spatial heterogeneity; detailed analysis of these differences shows that they arise mostly from a combination of improvements to the modelling framework. Average BII does not strongly differ between islands and mainlands, but richness-based BII has fallen by more on islands. It seems native species are more negatively affected by rising human population density and road development on islands than mainlands, and islands have seen more land conversion. Our results highlight the parlous state of biodiversity native to islands.

## INTRODUCTION

Biodiversity is continuing to decline and the pressures driving the declines are not easing (Butchart et al., 2010; Tittensor et al., 2014). The loss of biodiversity from ecosystems can compromise their functioning (Hooper et al., 2012) and therefore, their capacity to contribute to human wellbeing (Diaz et al. 2018). Ongoing debates have tried to identify a set of suitable indicators to inform about the state of biodiversity and progress towards biodiversity targets such as the Aichi 2020 Targets. The Convention on Biological Diversity (CBD) established a set of criteria that indicators of biodiversity change should satisfy; for example, indicators should be sensitive to changes at policy-relevant spatial and temporal scales, be easily aggregated and disaggregated to any spatial scale, allow comparisons with a baseline situation and be affordable and simple to understand (CBD, 2003; Scholes and Biggs, 2005).

The Biodiversity Intactness Index (BII) was proposed by Scholes & Biggs (2005) with these criteria in mind. BII was defined as “the average abundance of a large and diverse set of organisms in a given geographical area, relative to their reference populations”. The index provides estimates for biodiversity loss as a result of human pressures by focusing in the status of originally present species in a reference condition, which is represented by minimally-disturbed sites since historical data are very rare (Scholes and Biggs, 2005). Decreases in BII – i.e., the loss or population decline of originally present species – may capture falling ecosystem resilience and ecosystem’s ability to continue to meet societal needs: e.g., when facing disturbances, a high diversity ensures the persistence of at least a few species which might continue delivering ecosystem services – Biggs et al., 2012). BII initially only tested the effects of land use-among the main drivers of biodiversity loss (Maxwell et al. 2016; Brummitt et al. 2015) – without taking into account other related pressures such as human population growth.

BII has been proposed as a metric for assessing biotic integrity in the Planetary Boundaries framework (Steffen et al., 2015), which addresses the effects of human pressures on global sustainability and defines a safe operating space for humanity (Rockstrom et al., 2009, Steffen et al., 2015). The Planetary Boundaries framework places a safe limit at 10% reduction of BII, but this limit is highly uncertain; a less conservative estimate lies at 70% reduction (Steffen et al., 2015). The proposed safe limit focuses on the diversity needed within ecosystems to ensure the long term maintenance of ecosystem function (Mace et al. 2014, Steffen et al., 2015).

BII began as an index that relied on expert opinion and focused on specific geographical areas instead of global analyses, because of the lack of empirical data (Scholes and Biggs, 2005). The PREDICTS (Projecting Responses of Ecological Diversity In Changing Terrestrial Systems) project has developed a new implementation of BII based on a global collation of site-level biodiversity in sites facing different land uses and related pressures. This implementation leads to the estimation of local BII, which can be averaged for any larger spatial scale (e.g., globally, by countries, biodiversity hotspots or biomes). This approach also permits the exploration of temporal changes in BII under real (De Palma et al. 2018) or possible future changes in land use and other pressures (Hill et al., 2018) to inform policy. Local BII estimates are therefore relevant for global biodiversity assessments such as those developed by the Group on Earth Observations Biodiversity Observation Network (GEOBON) and the Intergovernmental Science-Policy Platform on Biodiversity and Ecosystem Services (IPBES). The attributes of the BII also link the index to the Essential Biodiversity Variables (EBVs) framework from GEOBON, since in addition to contributing to develop biodiversity forecasts under different policy and management scenarios, BII is a global biodiversity change indicator which is sustainable, technically and economically feasible, can link monitoring initiatives and decision makers, integrate remote sensing with in-situ observations and contribute to the assessment of progress towards the 2020 Targets of the CBD.

Recently, the PREDICTS project produced global BII estimates, finding that land use and related pressures have reduced local biodiversity intactness below the planetary boundary across 58% of the world’s land surface (Newbold et al., 2016). Newbold et al. (2016) warned about how previous data and models were not adequate to perform analyses for biomes and taxa separately, and pointed out that estimates could be biased for certain systems. Therefore, they did not take into account differences in responses in different ecological systems, such as islands and mainlands – two systems that differ greatly in terms of environmental conditions, species assemblages and human pressures; which can result in a different magnitude in the reduction of biodiversity intactness caused by land use change.

Islands might be suffering a greater decline of native species than mainlands. Islands’ small size can facilitate access to undisturbed areas, promoting a fast acceleration of habitat loss (Kier, 2009), whereas islands’ isolation can prevent the recovery of declining populations since it drives low rates of natural immigration (Lomolino, 1986). Moreover, island endemic species usually lack the potential to face disturbances in their habitat since evolutionary isolation often leads to the loss of traits that ease populations’ recovery after disturbances; for example, some island species present low reproductive rates (Sakai et al., 1995) and poor dispersal abilities (Gillespie et al., 2008). Islands are also more vulnerable to the establishment of alien species than mainlands (Sax & Brown, 2000, Whittaker & Fernández-Palacios, 2007), perhaps related to their species poverty (a result of island isolation and low colonisation rates -Whittaker & Fernández-Palacios, 2007) or high resource availability (Denslow, 2003). The introduction of alien species has led to extreme declines of island native populations (Whittaker & Fernández-Palacios, 2007). The decline of biodiversity intactness on islands is especially alarming since it might involve the loss of a large number of endemic species with unique evolutionary histories (Whittaker & Fernández-Palacios, 2007).

Here we update the BII estimates from Newbold et al. (2016), allowing island and mainland assemblages to respond differently to drivers. We test whether biodiversity intactness has decreased more severely on islands than mainlands as a result of larger decreases of local diversity and larger changes in species composition in human-dominated land uses. We also explore the intensity of land use and related pressures on islands and mainlands to determine whether this factor contributes to differences among biodiversity intactness of these ecological systems.

We calculate BII for islands and mainlands based on models focusing on how land use, human population density and distance to roads affect local biodiversity and models of how land use affects the similarity of species assemblages to assemblages in minimally-disturbed sites. We use estimates based on abundance and richness data and generate fine-scale (∼1km2) global maps for BII estimates that address some previous criticisms of BII (see Rouget et al., 2006 and Faith et al., 2008). These estimates differ somewhat from those published originally; we use a series of model comparisons to quantify the contribution to these differences of more data, more sensitive modelling of compositional similarity, and permitting different responses on islands and mainlands. In the process, we update the methods from Newbold et al. (2016), creating an approach that is suitable for analyses focusing on smaller datasets, which contributes to future analyses of BII for different biomes or clades.

## METHODS

All the models used data on species abundance and occurrence extracted in October 2016 from the PREDICTS database (Hudson et al. 2017). The database collates data from published research that compared local biodiversity across sites facing different land uses and related pressures. The PREDICTS database is structured hierarchically into Data Sources (publications), Studies (sampling method within a source), Blocks (spatial blocks, if present in the study) and Sites (Hudson et al., 2017).

We classified each site in the database as an island or mainland site by matching site coordinates with a global layer of land polygons taken from OpenStreetMap (OpenStreetMap Contributors, 2015). Australia and all land polygons with a smaller area were classified as islands. We classified Australia as an island since many of its characteristics are more island-like than continental: e.g., relatively small size, complete isolation from other continents by ocean and long isolation history (complete isolation ∼ 33 Mya -Wilford & Brown, 1994). Treating Australia as an island also helped to improve the balance between island and mainland sites in our analyses (though we tested the influence of this decision on our findings; see below).

### Statistical modelling

#### Biodiversity Intactness Index

Our approach for estimating BII requires combining two statistical models: a model of how land use and related pressures affect overall abundance or species richness at a site, and a model of how land use affects compositional similarity of assemblages to baseline assemblages (minimally-disturbed primary vegetation). Originally, Scholes & Biggs (2005) explicitly excluded alien species from the calculation of BII; however, since it is usually difficult to classify species into natives and aliens, the models of compositional similarity are used to calculate BII excluding species that are not present in minimally-disturbed primary vegetation (here referred as novel species).

BII calculation was performed based on species abundance following the original approach by Scholes & Biggs (2005); however, we also calculated BII using species richness to address previous critics – i.e., abundance-based BII might overlook species losses and calculate overoptimistic estimates if the abundance of the remaining species in disturbed sites increases (Faith et al., 2008). Using abundance-based and richness-based estimates also provides information about different consequences of BII decline for ecosystem functions and services. For example, abundance-based metrics give more weight to common species; therefore, the use of total abundance might be more appropriate when analysing the amount of ecosystem service provision. However, richness-based metrics give equal weight to rare and common species, which might be more relevant for stability of ecosystem service provision.

All statistical analyses were performed using R Version 3.2.3 (R Core Team, 2017). The analyses are based on Generalized mixed-effect models (GLMMs), which were fitted using the *lme4* package ver. 1.1-15 (Bates et al., 2017). We use GLMMs to deal with the heterogeneity among the studies’ methods, taxonomic focus and location (e.g., differences in sampling method and effort, sampled taxa and broad-scale biogeographic differences) and among spatial blocks (for studies with blocked or split-plot designs). Such differences might affect the estimation of site-level diversity; therefore, GLMMs provide a robust approach for quantifying the variation among studies (random-effects) without directly considering their effect in the analysis (Bolker et al., 2009). All the models fitted for BII estimation are adapted from Newbold et al. (2016).

##### 1. Modelling total abundance and species richness

As site-level measures of biodiversity, we calculated each site’s total abundance (sum of abundances of all present taxa) and species richness (number of unique present taxa). In cases where the sampling effort varied among sites in one study, total abundance was divided by the sampling effort to make data comparable among the study’s sites.

Sites in the database had previously been classified into 10 land-use categories and three land-use intensities (Minimal, Light and Intense) within each land use (details on all categories in Hudson et al. 2017). The ten land-use categories were collapsed into six final classes (primary vegetation, secondary vegetation, plantation forest, cropland, pasture and urban) to give reasonable sample sizes for both islands and mainlands. We created an additional site variable by combining land use and use intensity of each site (LUI). Sites were classified in 18 final LUI categories (Table S1). Primary vegetation with minimal use (here referred as PriMin) was considered as a baseline representing minimally disturbed sites.

We obtained site-level data for human population density (HPD) and distance to the nearest road (DistRd). We extracted data for these variables by matching site coordinates with global layers of human population density and the world’s roads. Human population density data (for the year 2000) was obtained from the Global Rural-Urban Mapping Project, Version 1 (GRUMPv1): Population Density (CIESIN, 2011). Data for distance to the nearest road was extracted from the Global Roads Open Access Data Set, Version 1 (gROADSv1) (CIESIN, 2013). Sites missing data for any of the human pressure variables were excluded from these models.

To model the responses of overall abundance on islands and mainlands, we fitted an initial maximal model where total abundance was analysed as a function of site-level land use and intensity category (LUI), human population density, distance to the nearest road and Island/Mainland classification (fixed-effects). We included two-way interactions between the Island/mainland term and the three human pressures plus three-way interactions between the Island/Mainland term, land use and HPD or DistRd. Prior to analysis, total abundance was rescaled to a zero-to-one scale within each study to reduce the variance among studies caused by differences in the taxonomic focus and sampling effort. The rescaled total abundance was then square-root transformed and modelled using a Gaussian error structure. We used square-root transformation since it resulted in a better residual distribution than log transformation (Figure S1). Models were fit with Gaussian errors rather than using untransformed data with Poisson errors because not all abundance data were integers (before rescaling within study), so a discrete error distribution like Poisson could not be used. HPD and DistRd were log transformed and then rescaled to a zero-to-one scale to deal with extreme values and to reduce collinearity. HPD and DistRd were fitted as quadratic orthogonal polynomials in the models.

Akaike’s Information Criterion (AIC) was used to determine the best random-effects structure for the maximal model fitted using Restricted Maximum Likelihood (REML). We assessed three possible random-effects structures; these varied in terms of the random slopes, but always included random intercepts of study and block within study to account for methodological differences among studies and the spatial structure of sites within studies. Apart from these random intercepts, the models that were compared included: (i) random slopes for land uses within study, (ii) random slopes for land uses + use intensity within study and (iii) no random slopes. The random slopes account for the variation among studies in the relationship of sampled biodiversity with land use and land-use intensity. The fixed-effects structure of the final model was determined using backwards stepwise model simplification with the model fitted using Maximum Likelihood.

We modelled species richness using the same methods as for the abundance model. In this case, the richness model was fitted using a Poisson error structure and log link and using Maximum Likelihood (ML). Species richness was not rescaled within each study. We tested for overdispersion in the richness model using the ‘*blmeco*’ package ver. 1.2 (Korner-Nievergelt et al., 2018). The dispersion measure computed by the ‘*dispersion_glmer*’ function suggested no overdispersion problem.

For both abundance and richness models, prior to modelling, we assessed multicollinearity for all explanatory variables using Generalized Variance Inflation Factors (GVIFs) (Zuur et al. 2009). All values for both models were below 3, indicating that there was no strong collinearity within the sets of explanatory variables (Zuur et al. 2009). Model diagnostics for the final abundance and richness models showed that both models fulfilled homogeneity and normality assumptions (Figures S1b and S2).

##### 2. Modelling abundance and richness-based compositional similarity

Additional GLMMs were fitted to estimate compositional similarity of assemblages in the different land uses to assemblages in minimally-disturbed primary vegetation on islands and mainlands. We modelled compositional similarities between pairs of sites as a function of the land use where both sites were located (i.e., land-use contrasts such as PriMin-Cropland), the geographic distance among sites, the environmental differences between them and the Island/Mainland location of the pair of sites. All studies in the PREDICTS database are either on a mainland or an island; therefore, no pairwise comparison was between an island site and a mainland site. For these models we excluded studies sampling only one species and studies where sampling effort varied among sites. Self-comparisons of sites were also removed, as well as pairs of sites where environmental distance could not be calculated due to missing data and where geographic distance equalled zero due to lack of coordinate precision for the sites.

As the response variable, we calculated compositional similarities between all possible pairwise comparisons within studies (making both forward and reverse comparisons between every pair of sites). Compositional similarity between sites was calculated based on total abundance and species richness, using asymmetric versions of the Jaccard Index (Newbold et al., 2016) (See Methods in Supplementary material). These asymmetric metrics are the most appropriate similarity measures since they reflect the proportion of the species or total abundance of a site that is made up of species also present in a minimally-disturbed site (Purvis et al., 2018).

For these models, land use and use intensity of the sites were collapsed into seven classes: 1) Primary vegetation with minimal use, 2) Primary vegetation (sites with light and intense use), 3) Secondary vegetation, 4) Plantation forest, 5) Cropland, 6) Pasture and 7) Urban (last five classes including all use intensities). The pairs of sites in the dataset had a total of 49 land-use contrasts (all possible combinations of the seven classes). However, to calculate BII we exclusively used the model estimates for compositional similarity between minimally-disturbed primary vegetation and the other six land uses (i.e., we only use and discuss results of six land-use contrasts). These estimates are the ones needed to correct the estimates of the abundance and richness models by excluding the proportion of species or total abundance that correspond to species that are not present in minimally-disturbed primary vegetation. We did not use LUI categories for the land-use contrasts since data were more limited for these analyses – i.e. only studies sampling PriMin can contribute to the calculation of compositional similarity of the land-use contrasts of interest–.

Geographic distance between the sites was included in the models to account for the distance-decay relationship (Nekola and White, 1999), and environmental distance to account for decay of similarity with environmental distance. To calculate environmental distance, we extracted site-level data for altitude and four bioclimatic variables at 1-km spatial resolution: maximum and minimum temperature and precipitation of wettest and driest month (Hijmans et al., 2005). We then calculated a single measure of environmental distance between each pair of sites was as the Gower (1971) dissimilarity between the five environmental variables, using the ‘*gower_dist*’ function in the ‘*gowe*r’ package ver. 0.1.2 (van der Loo, 2017). We estimated geographic distance for each pair of sites using the sites’ coordinates and the ‘distHaversine’ function in the ‘*geosphere*’ package ver. 1.5-7 (Hijmans et al., 2017).

We ran two separated models for abundance-based and richness-based compositional similarity. Fixed effects of both models included two-way interactions between the Island/Mainland term and the other three explanatory variables. The models included random intercepts of study. The PriMin-PriMin contrast was used as intercept condition in the models. This contrast reflects the natural spatial turnover of species, making it a natural baseline against which to compare the other land-use contrasts.

The compositional similarity data (ranging from 0 to 1) were logit-transformed prior to analysis using the ‘*car*’ package, ver. 2.1 (Fox & Weisberg, 2011), using a data adjustment of 0.01 to deal with 0 and 1 values (Warton et al., 2011). Logit transformation has advantages in power and interpretability over other transformations of proportional data (Warton et al., 2011). Environmental and geographic distances were also transformed, because both included extreme values (Zuur, et al., 2007). We tested different transformations and chose those where the data for each variable most closely approached a normal distribution. Environmental distance was transformed using cube root. Geographic distance was first divided by the median maximum linear extent of sites in the dataset and then log-transformed; this meant that a transformed geographic distance of zero corresponds to sites separated by their median linear dimension (i.e., adjacent sites). Model diagnostics suggest that our data treatment was adequate (Figure S9).

The final datasets for the compositional similarity models had extensive pseudo-replication because pairwise comparisons that involve the same sites are not independent. Standard statistical approaches could not be used to simplify the fixed effects of the models; therefore, we performed tests by permutation to address the dataset’s pseudo-replication and to determine whether fixed-effects of the models could be simplified without losing explanatory power. Our permutation tests are conceptually identical to the treatment of non-independence in multiple regression on distance matrices (MRM) (Lichstein, 2007), an extension of partial Mantel analysis (Smouse et al. 1986).

To perform backwards stepwise model simplification, we first performed normal likelihood ratio tests for our full model against the reduced model. As a second step, we permuted the model dataset by randomly shuffling the compositional similarity data within studies (i.e., rows of the response variable within each study) while holding all explanatory variables constant. Permutations were performed using the ‘*permute’* package ver. 0.9-4 (Simpson, 2013). We then ran the full and reduced models with the permuted dataset and performed a likelihood ratio test. We repeated this permutation process 199 times to generate null distributions for the likelihood ratios. This number of permutations was enough to get distributions approaching normality. p-values were calculated by performing a “greater” hypothesis test, where we compared the likelihood ratio from our models (observed value) against the distribution of the null likelihood ratios (comparison of 200 values: observed value + 199 null likelihood ratios). p-values were obtained using the ‘*as.randtest’* function from the ‘*ade4’* package ver. 1.7-10 (Dray et al., 2007), which compares the observed value to the null distributions by performing a Monte-Carlo test. Our test analysed whether the likelihood ratio of our models was higher than expected based on a model comparison with the same difference in degrees of freedom but no real loss of explanatory power. The test was performed only for the three interaction effects in our models since model simplification was not possible; i.e., all interaction effects showed significant p-values (0.005 in all cases: lowest value that can be obtained in our tests, which compare 200 values). We followed the same approach, to assess the statistical significance of the coefficients for the interactions between island/mainland and the other explanatory variables (Table S9).

The overall model intercepts estimated (logit-transformed) compositional similarity, for PriMin-PriMin contrasts on islands when (transformed) environmental and geographic distances were zero. Estimates for (logit-transformed) compositional similarity of each land-use contrast on islands were obtained by adding the model intercept to the contrast coefficients for islands; comparable values for mainlands were obtained by adding the model intercept and the PriMin-PriMin mainland coefficient to the contrast coefficients for mainlands plus each of the same island contrast coefficient. Calculating the inverse logit for these final island and mainland values converted them back to the original scale from 0 to 1. As a final step, the backtransformed values (on a scale from 0 to 1) for compositional similarity of island and mainland land-use contrasts were rescaled so that the contrast of PriMin against itself had a value of 1 on both islands and mainlands; this was done to avoid conflating natural spatial turnover with the effects of land use.

Only two of the island studies included the PriMin-Urban contrast, which is too few for random effects to be estimated reliably (Bolker et al. 2009). Therefore, we also estimated this coefficient indirectly, as the product of estimated Secondary-Urban and PriMin-Secondary compositional similarities (in a 0 to 1 scale) – as coefficients for these contrasts could be estimated from 10 and 38 island studies, respectively (enough for reliable estimation: Bolker et al. 2009). These indirect estimates were logit-transformed (using same data adjustment as before) and the respective model’s intercept (PriMin-PriMin island contrast) was subtracted to transform these values into model coefficients. Good agreement between the two estimates of the PriMin-Urban compositional similarity on islands would suggest that the original estimate is reasonable; but there was a strong disagreement (Table S9) which indicates the need for caution in interpreting it. Although based on more data, this second estimate makes the additional assumption that assemblages in secondary vegetation are directly between assemblages in primary minimal and urban sites on a straight line through multidimensional space; this assumption means that this second approach is likely to underestimate the true similarity between PriMin and Urban to some degree. Given the uncertainty around the original estimate, we have used the indirect estimate, but are careful not to place undue emphasis on this land-use contrast on islands.

##### 3. Calculation of BII and spatial projections

To calculate BII and perform global projections, we first separately map the modelled responses of overall abundance, species richness and compositional similarity (abundance-based and richness based). We used global pressure data at a resolution of 30 arc sec (∼1 km2) for each of the human pressure variables. We used the land use maps generated in Newbold et al., 2016, by downscaling (Hoskins et al., 2016) the harmonized land-use dataset for 2005 (Hurtt et al., 2011). No map was available for plantation forests since global land-use layers rarely distinguish this land use from other forests. Therefore, we modelled biodiversity responses to plantation forest but we omitted this effect when performing the global projections and therefore from BII calculation. Land-use intensity maps were generated using the statistical models in Newbold et al., 2015. Under this approach, the Global Land Systems dataset (van Asselen & Verburg, 2013) is reclassified into land-use and use-intensity combinations and models are fitted to estimate how the proportion of each 0.5 degree grid cell under each land use-intensity combination depends on the proportion of the grid cell under a particular land use, human population density and United Nations sub-region.

The gridded map for human population density and a vector map of the world’s roads (both for the year 2005) were obtained from NASA’S Socioeconomic Data and Applications Centre (CIESIN, 2011 and CIESIN, 2013). Following the methods by Newbold et al., 2016, we computed the gridded map of distance to nearest road using a Python code written for the arcpy module of ArcMap Version 10.3 (ESRI, 2011). The vector map was first projected onto an equal-area (Behrmann) projection to then calculate the average distance to the nearest road within each 782-m grid cell using the ‘*Euclidean Distance*’ function. Finally, the map was reprojected back to a WGS 1984 projection at 30 arc sec resolution.

The global maps of human pressures were separated into island and mainland maps (i.e. maps were clipped using island and mainland shapefiles derived from OpenStreetMap (OpenStreetMap Contributors, 2015). These maps were used to separately drive the four statistical models of how island and mainland biodiversity respond to pressures. For projections of the abundance and richness models, the data for HPD and DistRd in the maps were not allowed to exceed the maximum values in the modelled datasets to avoid extrapolations beyond our data (i.e., all grid cells with higher values than the maximum value in our data were set to this maximum value). The data in the HPD and DistRd maps were log-transformed and rescaled as in the models. For projections of total abundance and species richness, the values were back-transformed and expressed relative to minimally-disturbed primary vegetation with zero human population and at a distance to the nearest road equal to the maximum value in the final data used in the models (195.3 km). For compositional similarity projections (abundance-based and richness-based), we used models that included the indirect estimate for the PriMin-Urban contrast on islands. Compositional similarity values were back-transformed and expressed relative to compositional similarity among minimally-disturbed primary vegetation sites with zero environmental distance and zero geographic distance (in the latter case, this value equals to the median sampling extent in the dataset). BII for islands and mainlands (abundance-based and richness-based) was calculated by multiplying these spatial projections; e.g. island abundance-based BII is the product of island projections for responses of overall abundance and projections for abundance-based compositional similarity. We present our BII estimates in a 0 to 1 scale where 1 = 100% intactness.

The final BII maps for islands and mainlands shared 23,460 cells. These are cells that intersected with both the island and mainland shapefiles since islands smaller than 1km2 that are close to mainland coasts fit into cells that include mainlands (see Figure S10). Nevertheless, these cells were not a major concern since BII was calculated separately for islands and mainlands; therefore even though these cells share the same human pressure data, BII values in the island and mainland maps correspond to the modelled island or mainland responses. Moreover, the shared cells only represent 0.1% of the cells for the final island BII maps.

We calculated average BII values for island and mainland biomes (Olson et al., 2001), IPBES regions (Brooks et al., 2016), countries and the Conservation International’s biodiversity hotspots (Myers et al., 2000). We also calculated average BII values for islands listed in the Global Island Database, ver. 2.1 (UNEP-WCMC, 2015). Values were calculated by averaging modelled BII values across all cells intersecting the corresponding region shapefiles after reprojecting the BII maps and the shapefiles to a Behrmann equal-area projection. The intersection of the maps was performed using the separated island/mainland BII maps and separated island/mainland maps for countries and biomes (the global shapefiles were clipped using island and mainland shapefiles). This was done to ensure that calculations were performed exclusively for island or mainland regions considering the problem of shared cells on island and mainland BII maps. Not cutting the global shapefiles would have led to the inclusion of mainland regions in the island averages and vice versa. We used the global shapefiles for IPBES regions and hotspots for the intersection with the island and mainland BII maps since these regions are very broad. In the case of the shapefile from the Global Island Database, we added Australia as an additional polygon before intersecting it with the island BII maps. We calculated average BII for islands using the unique island ID code (id_gid) in the Global Island Database; averages for Australia were calculated using an exclusive code for the polygon.

Using the projected BII maps (Behrmann equal-area projection), we calculated the percentage of land surface on islands and mainlands that is below the recommended safe limit for reduction of biodiversity intactness: beyond 10% decrease –or 0.1 in our 0 to 1 scale – for abundance based BII (according to the proposed planetary boundary – Steffen et al., 2015) and beyond 20% decrease –or 0.2 in our scale– for richness-based BII (according to the limit used in Newbold et al., 2016, based on Hooper et al., 2012). Additionally, as a way to explore the intensity of human pressures on islands and mainlands that could drive differences among BII averages, we calculated the percentage of land surface under each land-use and use-intensity combination on islands and mainlands. We also explored the data distribution for HPD and DistRd on island and mainland maps. For these calculations we used projected (Behrmann equal-area) land-use intensity, HPD and DistRd maps for islands and mainlands, that only included cells that had a defined value in the final BII maps.

Since the classification of Australia as an island might be controversial, as a sensitivity test, we calculated island average BII values and the percentage of land surface under the safe limits using BII island maps where Australia (only the landmass considered as mainland Australia) was excluded. Considering that a high percentage of land surface in our original island maps corresponded to Australia, we also re calculated the percentage of land surface under each land-use and use-intensity combination using island use-intensity maps excluding Australia.

### Testing the effect of changes in BII modelling

Given that we have refined the modelling approach used by Newbold et al. (2016), we compared our model coefficients and BII estimates with theirs. We first compared estimates for the responses of total abundance and species richness to LUI. The main differences between our models and those in Newbold et al. are the inclusion of the island/mainland term and the data transformation for total abundance (in Newbold et al., total abundance was not rescaled to a zero-to-one scale within studies and total abundance was log-transformed). By plotting our island and mainland estimates against the global estimates from Newbold et al., we assessed whether previous global estimates were biased towards island or mainland data (i.e., which of the island or mainland estimates are more similar to the global estimates).

The compositional similarity models represent the main change in the methods for BII calculation. Previously, Newbold et al. used datasets only including independent pairs of sites within studies and averaged the coefficients of 100 models fitted with 100 different sets of pairwise comparisons that were randomly-chosen. This random selection led to a lower chance of picking up pairwise comparisons with land-use contrasts including land uses with fewer sites in the dataset. Or approach, using all possible pairwise comparisons within studies and performing permutation tests, allows us to use all available data in a single model. Further differences between the compositional similarity models are the baseline used in land-use contrasts (representing natural habitats) and the data transformation. Newbold et al. used primary vegetation (sites with the three use intensities) as baseline in the land-use contrasts, while we opted to only use minimally-disturbed primary vegetation as baseline for the contrasts. As for the data transformation, previously, all variables in the model were log-transformed, including compositional similarity data which is bounded between 0 and 1.

Using the abundance-based model, we tested whether using the new baseline and different data transformations (focusing on logit transformation for compositional similarity data) affected compositional similarity estimates. For these tests we used a subset of 30% of the pairwise comparisons of our abundance-based compositional similarity data (randomly sampling 30% of comparisons within each study); this percentage was enough to replicate our results and facilitated tests by reducing the time and computational power needed to run the models. To test the effects of logit transformation and the baseline on compositional similarity estimates, we first fitted our model log-transforming all the variables but keeping the original land-use contrasts to extract estimates of contrasts using PriMin and Primary vegetation (with light and intense use) as baseline. In a second test, before running the model, we collapsed land-use contrasts using PriMin and Primary vegetation to replicate the baseline used by Newbold et al.; we then ran a model using our data transformations and a second one log-transforming all the variables. Using the mainland estimates from each of these models, we calculated the average compositional similarity across all land-use contrasts (using rescaled values). We first compared the average for mainland land-use contrasts from our final abundance-based model against the average from the model replicating the conditions of models in Newbold et al. Next, based on averages from models with intermediate conditions (i.e. model with PriMin baseline + log transformation and model with Primary baseline + logit transformation) we defined whether the new baseline or the logit transformation was the main factor driving differences between our estimates and estimates generated by following methods from Newbold et al. (i.e., which average was closer to the average from our final model) (Table S13).

Finally, we checked differences between our final BII estimates for islands and mainlands and the global BII estimates from Newbold et al. We first mapped the differences between the estimates by subtracting our abundance and richness global BII maps (joining island and mainland maps – Figure 1) from the global maps from Newbold et al. With these maps we aimed to identify areas were BII estimates from Newbold et al. were lower, higher or very similar to ours. We also separated the global BII maps from Newbold et al. into island and mainland maps by clipping them using our island and mainland shapefiles. We then compared average BII estimates for islands and mainlands derived from Newbold et al. maps with our averages, which are the result of models accounting for differences between islands and mainlands. Finally, we compared our average BII values for biomes and biodiversity hotspots for islands and mainlands against the biomes and hotspots global averages.

**Figure 1.**
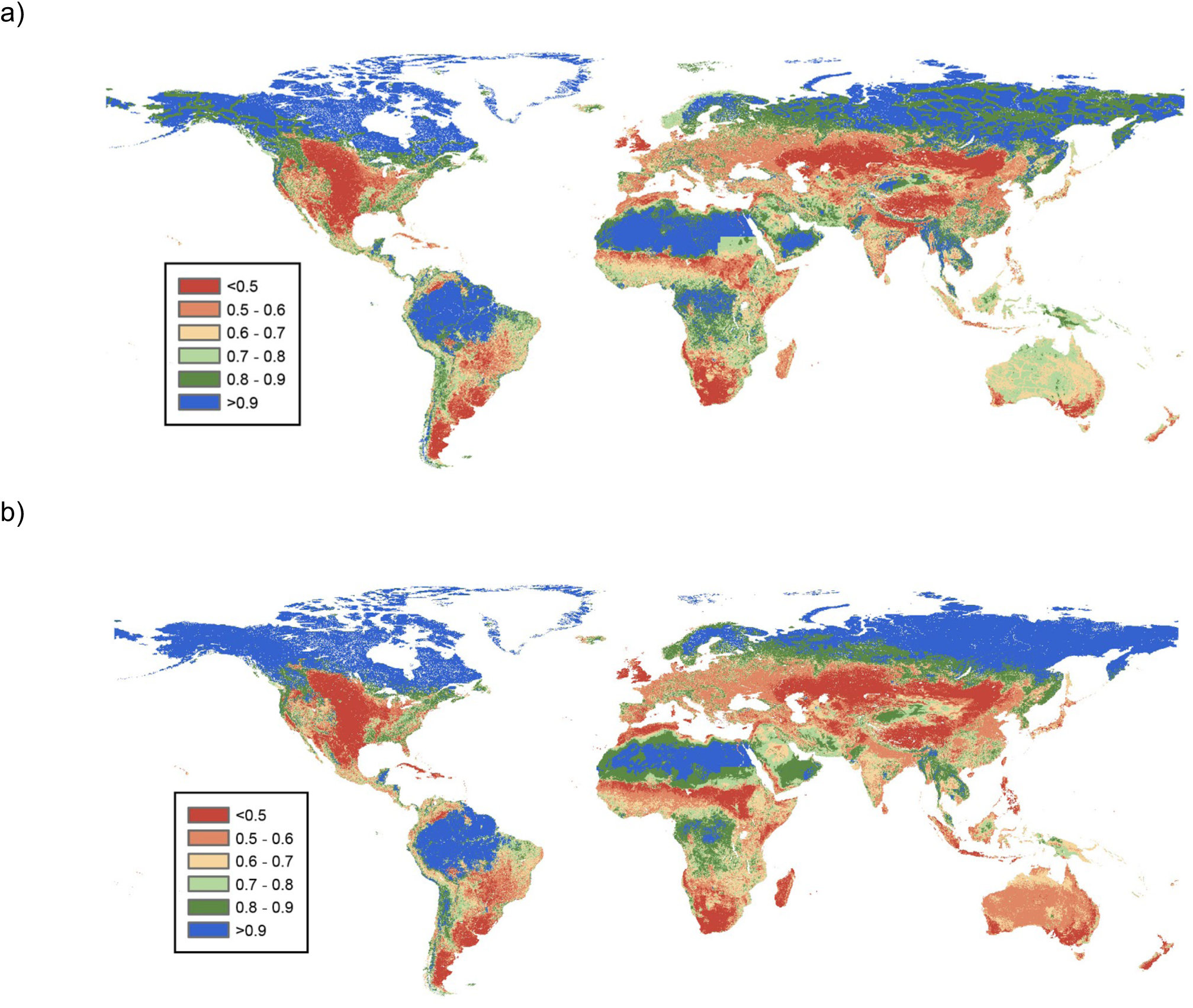
Global maps of biodiversity intactness. a) Abundance-based BII. b) Richness-based BII. BII values are shown in a 0 to 1 scale (1= 100% intactness)

**Figure 2.**
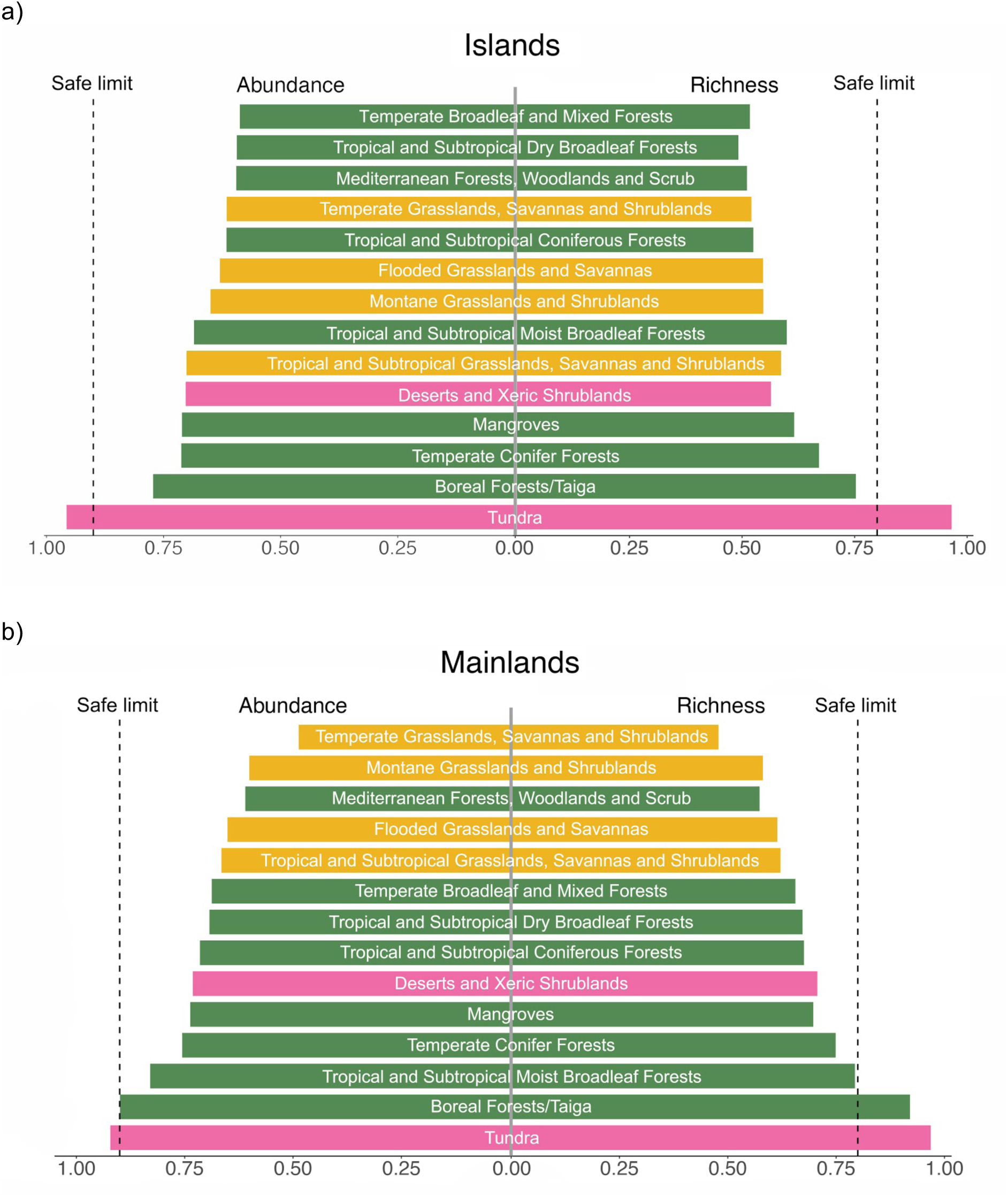
BII averages (abundance-based and richness-based) for the different biomes on a) islands and b) mainlands. The figures are replicas of those in Newbold et al. (2016) to ease comparisons. Colours indicate major biome type (green: forests, yellow: grasslands, pink: other).

## RESULTS

All the statistical models that were fitted for BII calculation suggested significant differences between island and mainland responses to human pressures. The abundance model was simplified since DistRd only influenced total abundance without interacting with any other variable (Figure S4). The rest of the interactions were kept in the model (Table S5), suggesting that the effects of LUI on total abundance differ between islands and mainlands (Figure S3) but also that total abundance is significantly influenced by interactions between land use, island/mainland location and HPD (Figure S5). The richness model retained all terms (Table S7), suggesting that the effects of LUI on species richness are different on islands and mainlands (Figure S6) and that species richness is significantly influenced by interactions between land use, island/mainland location and both HPD and DistRd (Figures S7 and S8). For both models the best random-effects structure included land use and use intensity as random slopes and random intercepts of study and block within study (Tables S4 and S6). However, due to convergence issues, in the final richness model we did not include use intensity as random slope. For some LUI (Figures S3 and S6) and particular combinations of human pressures (Figures S5, S7 and S8), the models suggested a steeper decline of total abundance and species richness on islands than on mainlands. However, this pattern was not clearly observed across combinations of pressures; in several cases, differences between island and mainland responses were less clear. In some cases, these models showed wide confidence intervals (e.g., Figure S3).

No model simplification was possible for models of compositional similarity (abundance-based and richness-based); all terms in the models were retained according to the permuted likelihood ratio tests (each interaction term was significant at p=0.005). The models suggested that when the effects of environmental differences and distance among sites are controlled for (Table S9), the effects of land use on compositional similarity to assemblages in PriMin differ significantly between islands and mainlands (Table S9). Based on the rescaled compositional similarity estimates, similarity of species assemblages in human-dominated land uses to assemblages in minimally-disturbed sites was lower on islands than on mainlands only in some cases (Table S10). For example, compared to mainlands, in the abundance-based models, islands showed lower compositional similarity to PriMin assemblages in primary and secondary vegetation and croplands, while for the richness-based model, island estimates were lower for compositional similarity between PriMin and primary vegetation, croplands and urban sites.

Our maps of BII (Figure 1) suggest that on average, BII is lower for islands than for mainlands based on both total abundance and species richness. Average abundance-based BII was 0.71 (s.d.= 0.13) for islands and 0.73 (s.d.=0.19) for mainlands. The difference between islands and mainlands was greater when BII was calculated based on species richness; in this case average BII was 0.62 (s.d.= 0.16) for islands and 0.71 (s.d.= 0.19) for mainlands. The standard deviations that we present for BII averages indicate how spatially variable BII estimates are; at a global scale, BII estimates were slightly more variable for mainlands than islands. Our results suggest that on both islands and mainlands, average local abundance and species richness of originally present species have fallen below the values proposed as safe limits (0.9 for abundance and 0.8 for richness). We estimated that around 88.6% of the land surface of islands is below the recommended abundance-based safe limit, while 85.9% of the land surface transgresses the richness-based safe limit. The percentage of mainland land surface below the safe limit was 76.5% when taking into account the abundance-based boundary and 59.7% based on the richness-based boundary.

Based on our land-use intensity maps for islands and mainlands, we found that the percentage of land surface with minimally-used primary vegetation is higher on islands than on mainlands (∼38% vs 28%) (Table 2). On the other hand, the percentage of land surface with pastures and urban sites (most use intensities – Table S16) was higher on islands than on mainlands. Mainlands showed a higher percentage of land surface with secondary vegetation and croplands (all use intensities) than islands (Table 2). In the case of HPD and DistRd data, island data was more skewed towards low values for HPD (Figure S11a) and high values for DistRd (Figure S11b) when compared to mailands.

When Australia was excluded from the island maps, the main changes were for percentages of land surface with minimally-used primary vegetation (increasing to ∼45%) and pastures (e.g., percentage of pastures with light use decreased from ∼30% to ∼10% – Table S16). Nevertheless, when excluding Australia from the island BII maps, there were no extreme changes for island BII averages even though they increased slightly (abundance-based BII= 0.73 and richness-based BII= 0.67). In the case of the land surface of islands below the recommended BII safe limits, when Australia was excluded, 77.11% of islands surface was below the abundance-based safe limit and 71.80 % of the surface was below the richness-based safe limit. These percentages were still higher than those found for mainlands.

Average BII varied among biomes on islands and mainlands (Figure 3). While on mainlands grasslands are most affected, on islands, various forest biomes are the most affected. Tundra and boreal forests are the least affected on both islands and mainlands. For islands, BII averages for all biomes except tundra are below the values proposed as safe limits (abundance-based and richness-based). In the case of mainlands, only tundra and boreal forests have BII averages within the safe limits. Average BII also varied among biodiversity hotspots on islands and mainlands. For islands, the Mediterranean Basin was the hotspot with the lowest BII average, followed closely mainly by hotspots in tropical realms (e.g., Coastal Forests of Eastern Africa, Caribbean Islands and Madagascar and the Indian Ocean Islands) (Figure 3a). The Tumbes-Chocó-Magdalena hotspot was the least affected for islands. In the case of mainlands, hotspots in temperate realms were most affected, being the Succulent Karoo hotspot the one with the lowest BII average. The Indo-Burma and Sundaland hotspots were the least affected for mainlands (Figure 3b). For both islands and mainlands, BII averages for all hotspots were below the safe limits (abundance-based and richness-based).

**Figure 3.**
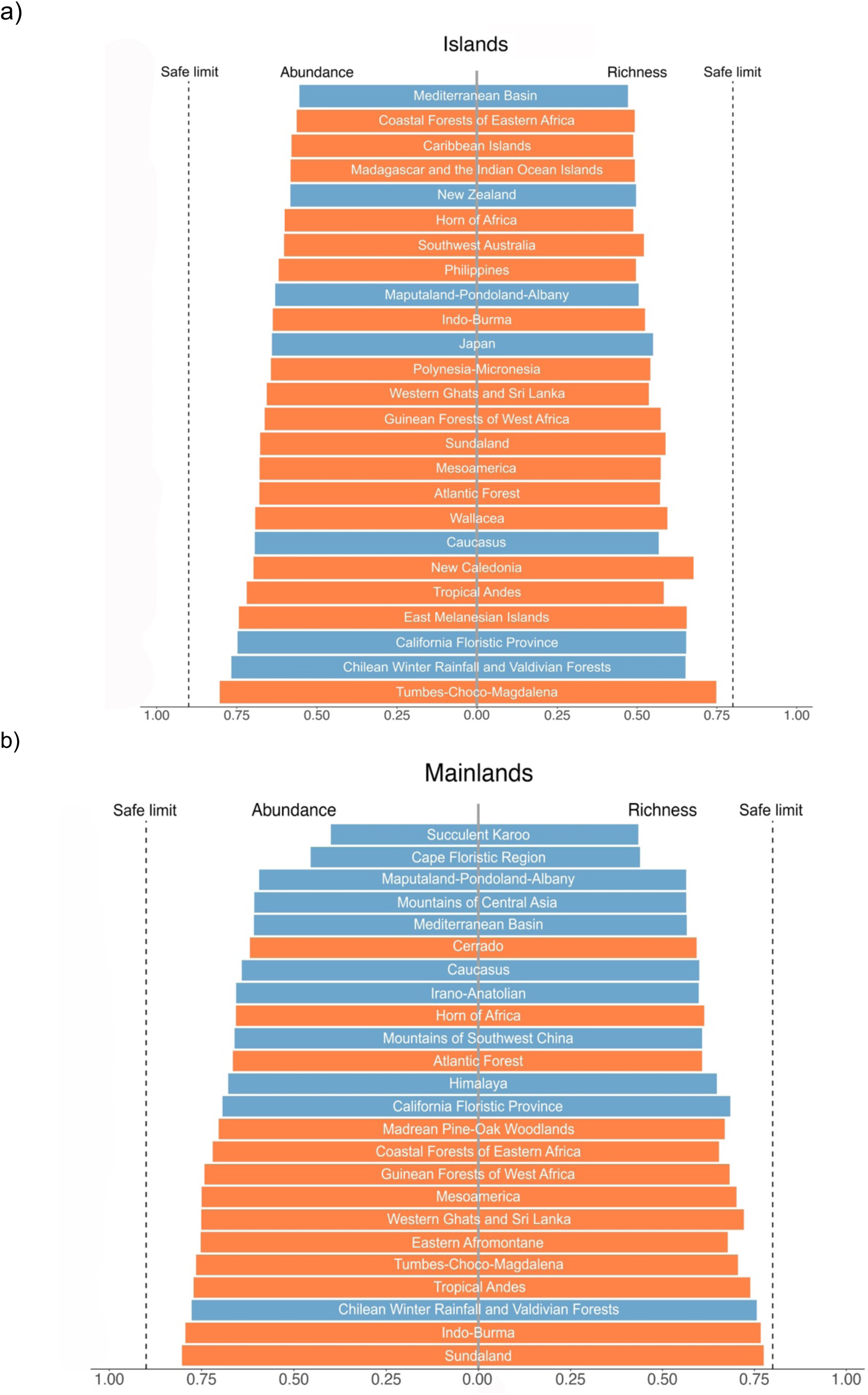
BII averages (abundance-based and richness-based) for the different biodiversity hotspots on a) islands and b) mainlands. The figures are replicas of those in Newbold et al. (2016) to ease comparisons. Colours indicate whether hotspots are in the tropical (orange) or temperate (blue) realms.

Values for average BII varied among countries (Figure 4); across regions, european countries had the lowest median value for average BII for both islands and mainlands. For islands, countries in Oceania (abundance-based) and Asia (richness-based) had the highest median values for average BII. For mainlands, the highest median value for average BII was for american countries. Overall, countries within each region had a lower median value for average BII for islands than mainlands (Figure 4). In the case of IPBES regions, average BII for Africa was the lowest for islands, while the Americas had the highest average for islands (abundance-based and richness-based) (Table 1). For mainlands, the lowest BII average was for the Asia-Pacific region, while averages for the other three regions did not differ considerably (abundance-based and richness-based) (Table 1). Once again, overall, all IPBES regions had lower BII averages for islands than for mainlands, with the only exception of the Americas (abundance-based and richness-based) and the Asia-Pacific region (abundance-based).

**Table 1.** Average BII (abundance-based and richness-based) for IPBES regions on islands and mainlands. s.d. are shown within parenthesis. Results for Areas Beyond National Jurisdiction (ABNJ) are not shown.

| IPBES region | Average BII abundance-based |  | Average BII richness-based |  |
| --- | --- | --- | --- | --- |
|  | Islands | Mainlands | Islands | Mainlands |
| Africa | 0.58 (0.06) | 0.75 (0.18) | 0.49 (0.05) | 0.70 (0.17) |
| Americas | 0.88 (0.14) | 0.75 (0.20) | 0.86 (0.18) | 0.74 (0.20) |
| Asia-Pacific | 0.68 (0.09) | 0.67 (0.17) | 0.57 (0.08) | 0.64 (0.15) |
| Europe-Central Asia | 0.75 (0.21) | 0.75 (0.18) | 0.72 (0.24) | 0.75 (0.20) |

**Table 2.** Percentage of land surface under different land-use and use-intensity combinations (LUI) in the island and mainland BII maps. LUI classes are collapsed as in the compositional similarity models. More detailed percentages (each land use with different use intensities) are shown in Table S16.

| LUI | Percentage on islands | Percentage on mainlands |
| --- | --- | --- |
| Primary Vegetation Minimal use | 38.45 | 28.48 |
| Primary Vegetation | 9.38 | 9.73 |
| Secondary Vegetation | 10.3 | 24.42 |
| Cropland | 8.98 | 12.29 |
| Pasture | 32.38 | 24.68 |
| Urban | 0.49 | 0.39 |

**Figure 4.**
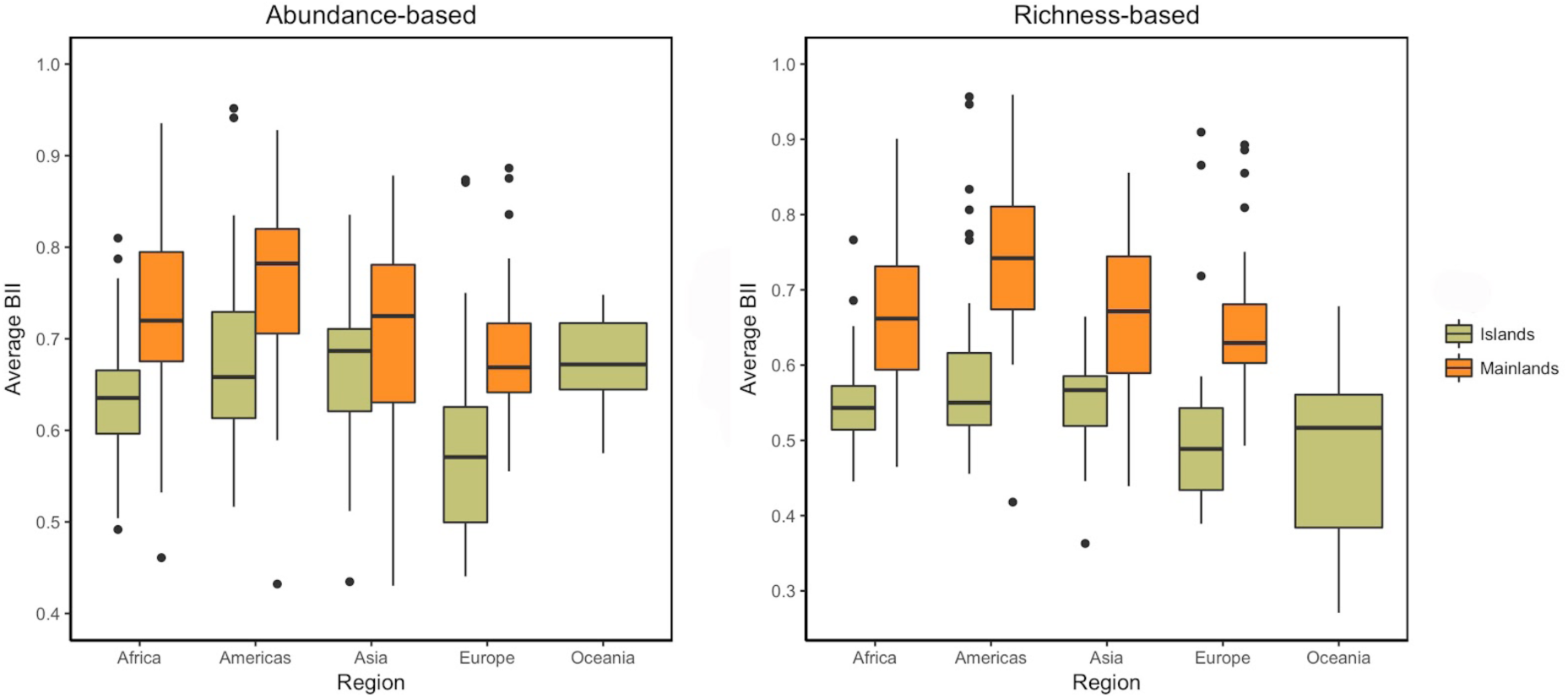
Average BII (abundance-based and richness-based) at the country level across different regions (UN geographic regions) on islands and mainlands. Lines within boxes show the median value, boxes show data within the 25th to 75th percentiles, whiskers show points that are up to 1.5 times the interquartile range of the data and points show countries that fall outside of these limits.

We were able to estimate BII averages for 3,602 islands (islands for which pressure data was available and whose name was listed in the Global Island Database) plus Australia. The island with the lowest BII average (for both abundance-based and richness-based) was Roosevelt Island in the U.S. with an average of ∼0.2 for BII. The islands with the highest BII averages (∼0.98) were mainly located in Greenland, Canada and Russia. BII averages for each island are publicly available at -----(doi: --------).

Our average BII values for islands and mainlands were lower than the global BII average calculated in Newbold et al. (2106), who reported ∼0.85 for abundance-based BII and ∼0.77 for richness-based BII. By mapping the differences between Newbold et al. estimates and ours, we found that our estimates are lower in most regions, with the main exceptions of Australia (abundance-based), the Scandinavian peninsula, Russia (richness-based) and the Sahara desert (abundance-based and richness-based) (Figure S18 a and b). BII estimates seemed to differ more extremely on islands, North America and South Africa (abundance-based and richness-based). Our abundance-based BII estimates were particularly lower than estimates from Newbold et al. In this case, a big proportion of Europe and Asia showed big differences between the estimates (Figure S18a).

Our averages were also considerably lower than the island and mainland averages obtained from the global BII maps (abundance-based and richness-based). When separating the global BII maps from Newbold et al. into island and mainland maps, for abundance-based BII, average for islands was 0.79 (s.d.=0.16), while for mainlands average was 0.85 (s.d.=0.13). Richness-based BII averages estimated from the global maps were 0.76 (s.d.=0.11) for islands and 0.77 (s.d.=0.10) for mainlands.

In the case of biomes, most of our average BII values for islands and mailands were lower than global averages for biomes from Newbold et al. (Figure S14 and S15). The only values that were higher than the global averages were richness-based averages for tundra (for islands and mainlands) and boreal forests (for islands) (Figure S15). Mainland averages for biomes were more similar to the global averages (especially in the case of richness-based BII – Figure S15), which resulted in the same pattern found in Newbold et al., where the lowest BII averages were found for grasslands (Figure 3). Island averages for biomes showed bigger differences to the global averages, especially for richness-based BII (Figure S15). All hotspots BII averages for islands and mainlands were lower than global averages estimated in Newbold et al. (abundance-based and richness-based) (Figures S16 and S17). Once again, averages for islands showed bigger differences to global averages when compared to mainlands. Mainlands showed the same pattern found in Newbold et al., in terms of the hotspots with the lowest BII averages (Figure 3b).

More particularly, for each of the statistical models that we fitted, we also found marked differences between our results and results from models in Newbold et al. In the case of the effects of LUI on total abundance and species richness, when comparing the global estimates from Newbold et al. against our island and mainland estimates, we found that results for the effects of some LUI were very similar for mainlands and the global estimates (Figures S12 and S13). Island results for the effects of some LUI showed slightly more variation from the global estimates (Figures S12 and S13). Estimates for responses of total abundance (Figure S12) for both islands and mainlands showed much bigger differences from the global estimates than estimates for richness responses (Figure S13). There was a tendency for our abundance estimates to be lower than those in Newbold et al. (Figure S12)

Our estimates for compositional similarity of assemblages in different land uses to assemblages in natural habitats (Table S10) were much lower (for both islands and mainlands) and showed a wider range than estimates from Newbold et al. Our tests showed that these differences are driven by both the logit transformation for compositional similarity data and the use of minimally-disturbed primary vegetation as a baseline representing undisturbed habitats. When fitting our compositional similarity model (abundance-based) with different conditions, we found the highest compositional similarity values in models using log transformation and in estimates for land-use contrasts using a baseline of collapsed primary vegetation (Tables S11 and S12). However, based on averages of compositional similarity of land-use contrasts from models with different conditions (Table S13), we found that the logit transformation is the main factor driving the differences from estimates in Newbold et al. We concluded this since the compositional similarity average from the model using logit transformation but a baseline of collapsed primary vegetation was the most similar to the average from our final model (Table S13).

## DISCUSSION

The Biodiversity Intactness Index (BII) has attracted increasing attention since being proposed as a suitable metric to assess biodiversity loss relative to the safe limit proposed by the Planetary Boundaries framework (Steffen et al. 2015). Recently, BII averages have been estimated using global models (Newbold et al., 2016, Hill et al., 2018), in some cases focusing on specific biomes (e.g., tropical and subtropical forests – De Palma et al., 2018). Those models have not accounted for differences between responses to human pressures in different ecological systems. Our comparison of BII on islands and mainlands is based on statistical models that, for the first time, allow assemblages in these ecologically distinct contexts to respond differently to human drivers and to have different distance-decay relationships of compositional similarity.

The proposed safe limit for biosphere integrity is an abundance-based BII of 0.9, though the uncertainty around how BII relates to large-scale ecosystem functioning (Mace et al., 2014, Steffen et al. 2015) is reflected by a wide range of uncertainty in where the threshold should lie (between 0.3 and 0.9: Steffen et al. 2015). As well as abundance-based BII, Newbold et al. (2016) also assessed richness-based BII against a possible threshold of 0.8, as the loss of 20% (0.2) of species often drives ecosystem change (Hooper et al. 2012). Our results suggest that on both islands and mainlands, average local abundance and richness of originally present species have fallen from their values in minimally-disturbed habitats to levels that may compromise ecosystem function (Hooper et al., 2012; Steffen et al., 2015).

Islands have lower BII values than mainlands for both abundance-based BII (0.71 vs 0.73) and richness-based BII (0.62 vs 0.71, though the island value rises to 0.67 if Australia is excluded). These differences may reflect both greater sensitivity and greater exposure of island assemblages to human pressures. Although the effect sizes for many land uses are similar between island and mainland models of abundance, richness and compositional similarity (Figures S3-S8; Tables S9-S10;), increasing human population density and decreasing distance to roads both have markedly negative effects on species richness in primary vegetation on islands but not on mainlands (Figures S7, S8). Human population density also reduces total abundance in primary vegetation and pastures more on islands than on mainlands, although these declines are not as steep as those for species richness in primary vegetation. In terms of exposure, although islands have a higher percentage of minimally-used primary vegetation than mainlands, they have also seen more widespread conversion of land to human use; e.g. 42% of land on islands is cropland, pasture or urban, compared with 37% on mainlands (Table 2). Ramankutty et al. (2008) suggested that some island-rich regions, such as Southeast Asia and the Pacific, have a high percentage of croplands and pastures. These, together with urban areas, are the land uses that usually involve the most severe changes from natural habitats (Foley et al., 2005; McKinney, 2006), and that were associated with the biggest decreases in our models of total abundance, species richness and compositional similarity. A significantly higher human pressure on islands than on mainland regions has previously been suggested by Kier et al., (2009), who used the Human Impact Index (Sanderson et al., 2002), as measure of current threat (largely referring to the year 2000). Islands’ small size can facilitate the access to remote areas with remaining primary vegetation (Kier et al., 2009) and brings a higher human population density in close proximity to natural habitats (Delgado et al., 2017), making islands more vulnerable to habitat loss. The steep declines in local island (but not mainland) richness as human population increases and proximity to roads decreases suggest that species native to islands may have greater need of ‘people-free space’ than those on mainlands; while the widespread land conversion on islands suggests that many do not get it.

The BII averages within countries and IPBES regions are consistently lower for islands than mainlands in most regions (Figure 4; Table 1); the main exception was the Americas (IPBES region), where average BII values were higher for islands than mainlands perhaps mostly because of remote high-latitude islands (Figure1). Average BII is below the proposed safe limits on both islands and mainlands, for all IPBES regions (apart from islands in the Americas using richness-based BII), all biomes except for tundra and boreal forest, and for all biodiversity hotspots. These results are much less optimistic than those of Newbold et al. (2016), who estimated that only 9 of the 14 terrestrial biomes and 22 of the 34 terrestrial biodiversity hotspots have on average transgressed the safe limits for BII.

Our BII averages for mainlands and especially islands are much lower than those from Newbold et al. (2016). Our lower averages for abundance-based BII might be partially driven by our abundance model estimating bigger decreases of total abundance in some LUI classes – for both islands and mainlands – than in Newbold et al.’s (2016) global model. Our models are based on a larger volume of data and we also rescaled total abundance within each study to have a maximum value of 1, to reduce the otherwise high among-study heterogeneity. However, the main reason our BII estimates are lower is that our estimates of compositional similarity are lower. Whereas Newbold et al.’s (2016) estimates for similarity between other land uses and primary vegetation ranged between 0.90 and 0.99 (both abundance-based and richness-based similarity), our estimates ranged as low ∼0.6 for abundance-based similarity and ∼0.5 for richness-based similarity, for both islands and mainlands (see Table S10). These lower estimates of similarity arise partly from using minimally-disturbed primary vegetation as a baseline for sites’ comparisons (whereas Newbold et al. (2016) included all primary vegetation sites in the baseline) but mainly from the logit transformation of the compositional similarity data, which adequately reflects the data distribution and captures differences among natural habitats and other land uses better than the log-transformation used by Newbold et al (2016) (Table S13).

Separating the global maps from Newbold et al. (2016) into island and mainland also yields a lower BII average for islands than mainlands; however, islands stand out on ‘difference maps’ that compare our BII estimates with the earlier ones (Figure S18). This, together with the statistically highly significant differences between many island and mainland model coefficients, suggests that previous global estimates may have been biased towards the mainland picture, as the PREDICTS database has many more mainland than island studies. Not allowing island assemblages to respond differently from mainland assemblages appears to have led to optimistic estimates of BII for the more vulnerable systems, for which data is less commonly available.

Our study shares some limitations with other implementations of BII that have been discussed previously (Newbold et al., 2016, Purvis et al., 2018, De Palma et al. 2018). To summarise, our approach might be overestimating BII as a result of 1) ignoring lagged responses; 2) not considering the effects of climate change or other drivers that could have a different spatial pattern than the human pressures in our models; 3) the likely presence of human pressures on many sites in the reference class (assumed as minimally-disturbed); and 4) shortcomings in pressure data for some regions (e.g., lack of plantation forests in land-use maps, and bias and incompleteness in currently available road maps such as gROADS – Meijer et al., 2018). Effects of roads on islands might be underestimated if island roads are less likely to be recorded than those on mainlands. A further limitation more specific to this study is that many more studies are available for mainlands than islands, particularly within some biomes (see Tables S1, S2 and S8). Less confidence can also be placed in results for biomes that are relatively poorly represented in our data such as taiga, tropical and subtropical coniferous forests in islands; and tundra, mangroves and flooded grassland and savanna in both islands and mainlands (Table S2).

Despite these limitations, our study marks an important technical improvement in the estimation of BII, and shows that biotic integrity worldwide has been much more seriously diminished-especially on islands-than previously estimated (Newbold et al. 2016). Our results suggest that island biotas have lost more integrity partly because they may be more intrinsically sensitive and partly because they are more exposed to some human pressures. Species native to islands appear much less able to persist in the face of rising human population density and road development than mainland species. Our results highlight the need for stronger efforts to arrest the loss of native island biodiversity in order to achieve biodiversity targets, especially in the endemic-rich and heavily impacted island-based biodiversity hotspots (Myers et al. 2000).

