## Supplemental tables and figures for "Land-use and related pressures have reduced biotic integrity more on islands than on mainlands"

**SUPPLEMENTARY MATERIAL**

### Methods

#### *Calculation of compositional similarity*

We calculated two measures of compositional similarity between sites *i* and *j*: the richness-based asymmetric Jaccard Index, $J_R$ (= $S_{ij}$ /$S_j$, where $S_{ij}$ is the number of species common to both sites and $S_j$ is the number of species in site *j* – Newbold et al., 2016), and the abundance-based asymmetric Jaccard index, $J_A$ (= $A_{ij}$ /$A_j$, where $A_{ij}$ is the summed abundance at site *j* of all species common to both sites and $A_j$ is the summed abundance of all species at site *j* – Chao et al., 2005).

Table S1. Number of sites on islands and mainlands per LUI category in the final datasets for the abundance and richness models. The number of sites for the richness and abundance datasets differ since not all studies provide abundance data. Numbers in brackets show the number of studies from which data came from.

| LUI | Abundance dataset | | Richness dataset | |
|---|---|---|---|---|
| | Islands | Mainlands | Islands | Mainlands |
| Primary Vegetation Minimal use | 1012 (85) | 3299 (212) | 1370 (100) | 3778 (245) |
| Primary Vegetation Light use | 467 (38) | 1486 (108) | 520 (43) | 2026 (130) |
| Primary Vegetation Intense use | 252 (9) | 295 (35) | 266 (12) | 351 (41) |
| Secondary Vegetation Minimal use | 801 (82) | 1792 (166) | 944 (90) | 2200 (185) |
| Secondary Vegetation Light use | 633 (47) | 772 (90) | 715 (54) | 985 (97) |
| Secondary Vegetation Intense use | 393 (23) | 285 (41) | 414 (28) | 314 (44) |
| Pasture Minimal use | 911 (24) | 666 (53) | 911 (24) | 673 (57) |
| Pasture Light use | 1165 (38) | 1074 (66) | 1173 (40) | 1104 (72) |
| Pasture Intense use | 228 (22) | 1226 (22) | 241 (24) | 1232 (23) |
| Plantation forest Minimal use | 133 (20) | 354 (52) | 198 (24) | 507 (61) |
| Plantation forest Light use | 664 (30) | 686 (51) | 714 (36) | 722 (55) |
| Plantation forest Intense use | 76 (10) | 377 (28) | 90 (12) | 389 (31) |
| Cropland Minimal use | 49 (18) | 622 (50) | 51 (20) | 659 (54) |
| Cropland Light use | 78 (11) | 1111 (65) | 78 (11) | 1291 (69) |
| Cropland Intense use | 291 (17) | 1077 (45) | 292 (18) | 1176 (48) |
| Urban Minimal use | 147 (18) | 218 (20) | 149 (19) | 251 (27) |
| Urban Light use | 412 (21) | 221 (26) | 428 (22) | 273 (33) |
| Urban Intense use | 74 (8) | 38 (13) | 74 (8) | 196 (20) |

Table S2. Number of island and mainland sites per biome in the final datasets for the abundance and richness models.

| Biome | Abundance dataset | | Richness dataset | |
|---|---|---|---|---|
| | Islands | Mainlands | Islands | Mainlands |
| Boreal Forests/Taiga | 0 | 1049 | 0 | 1075 |
| Deserts & Xeric Shrublands | 55 | 210 | 55 | 218 |
| Flooded Grasslands & Savannas | 0 | 39 | 0 | 51 |
| Mangroves | 6 | 24 | 6 | 24 |
| Mediterranean Forests, Woodlands & Scrub | 303 | 1422 | 304 | 1614 |
| Montane Grasslands & Shrublands | 485 | 277 | 485 | 488 |
| Temperate Broadleaf & Mixed Forests | 3764 | 6790 | 4051 | 7607 |
| Temperate Conifer Forests | 98 | 422 | 98 | 512 |
| Temperate Grasslands, Savannas & Shrublands | 280 | 271 | 456 | 704 |
| Tropical & Subtropical Coniferous Forests | 0 | 277 | 0 | 367 |
| Tropical & Subtropical Dry Broadleaf Forests | 82 | 400 | 82 | 430 |
| Tropical & Subtropical Grasslands, Savannas & Shrublands | 545 | 1386 | 571 | 1721 |
| Tropical & Subtropical Moist Broadleaf Forests | 2168 | 3011 | 2520 | 3266 |
| Tundra | 0 | 21 | 0 | 50 |

Table S3. Numbers of species by major taxonomic group represented in the final datasets for the abundance and richness models.

| Taxon | Abundance dataset | | Richness dataset | |
|---|---|---|---|---|
| | Islands | Mainlands | Islands | Mainlands |
| | Vertebrates and invertebrates | | | |
| Amphibia | 136 | 291 | 136 | 317 |
| Annelida | 160 | 136 | 160 | 136 |
| Arachnida | 973 | 2554 | 974 | 2555 |
| Archaeognatha | 9 | 4 | 9 | 4 |
| Aves | 1012 | 3383 | 1077 | 3792 |
| Blattodea | 37 | 19 | 40 | 19 |

| | | | | |
|---|---|---|---|---|
| Chilopoda | 56 | 46 | 56 | 46 |
| Coleoptera | 2689 | 4092 | 2784 | 4094 |
| Collembola | 113 | 130 | 113 | 193 |
| Dermaptera | 9 | 10 | 9 | 10 |
| Diplopoda | 87 | 72 | 87 | 72 |
| Diplura | 2 | 1 | 2 | 1 |
| Diptera | 410 | 1128 | 521 | 1129 |
| Embioptera | 2 | 2 | 2 | 2 |
| Ephemeroptera | 2 | 2 | 2 | 2 |
| Hemiptera | 837 | 713 | 878 | 713 |
| Hymenoptera | 1349 | 3715 | 1654 | 3809 |
| Isoptera | 17 | 124 | 17 | 124 |
| Lepidoptera | 932 | 2881 | 1045 | 3197 |
| Malacostraca | 54 | 47 | 54 | 47 |
| Mammalia | 216 | 427 | 225 | 463 |
| Mantodea | 5 | 25 | 5 | 25 |
| Maxillopoda | 1 | 0 | 1 | 0 |
| Mecoptera | 1 | 2 | 1 | 2 |
| Megaloptera | 1 | 0 | 0 | 1 |
| Mollusca | 195 | 114 | 196 | 205 |
| Nematoda | 380 | 386 | 380 | 386 |
| Neuroptera | 14 | 34 | 17 | 34 |
| Odonata | 91 | 4 | 92 | 4 |
| Onychophora | 3 | 0 | 3 | 0 |
| Orthoptera | 37 | 152 | 99 | 170 |
| Pauropoda | 2 | 2 | 2 | 2 |
| Phasmida | 1 | 1 | 1 | 1 |
| Phthiraptera | 1 | 1 | 1 | 1 |
| Platyhelminthes | 4 | 1 | 5 | 1 |
| Plecoptera | 6 | 0 | 6 | 0 |

| | | | | |
|---|---|---|---|---|
| Protura | 1 | 1 | 1 | 1 |
| Psocodea | 33 | 3 | 33 | 3 |
| Raphidioptera | 0 | 1 | 0 | 1 |
| Reptilia | 183 | 239 | 197 | 243 |
| Siphonaptera | 1 | 1 | 1 | 1 |
| Symphyla | 5 | 1 | 5 | 1 |
| Thysanoptera | 25 | 18 | 25 | 18 |
| Trichoptera | 5 | 6 | 5 | 6 |
| Zoraptera | 1 | 0 | 1 | 0 |
| Zygentoma | 1 | 1 | 2 | 1 |
| | Plants | | | |
| Bryophyta | 271 | 918 | 302 | 1139 |
| Cycadopsida | 0 | 2 | 0 | 2 |
| Equisetopsida | 3 | 1 | 4 | 6 |
| Gnetopsida | 13 | 5 | 13 | 5 |
| Liliopsida | 785 | 1348 | 993 | 1623 |
| Lycopodiopsida | 16 | 9 | 18 | 11 |
| Magnoliopsida | 2853 | 5825 | 4053 | 7479 |
| Marattiopsida | 1 | 1 | 1 | 1 |
| Pinopsida | 12 | 33 | 23 | 45 |
| Polypodiopsida | 208 | 125 | 234 | 170 |
| Psilotopsida | 8 | 1 | 10 | 1 |
| | Fungi | | | |
| Ascomycota | 469 | 340 | 470 | 400 |
| Basidiomycota | 413 | 369 | 413 | 519 |
| Glomeromycota | 20 | 11 | 20 | 11 |
| | Protozoans | | | |
| Mycetozoa | 2 | 1 | 2 | 1 |

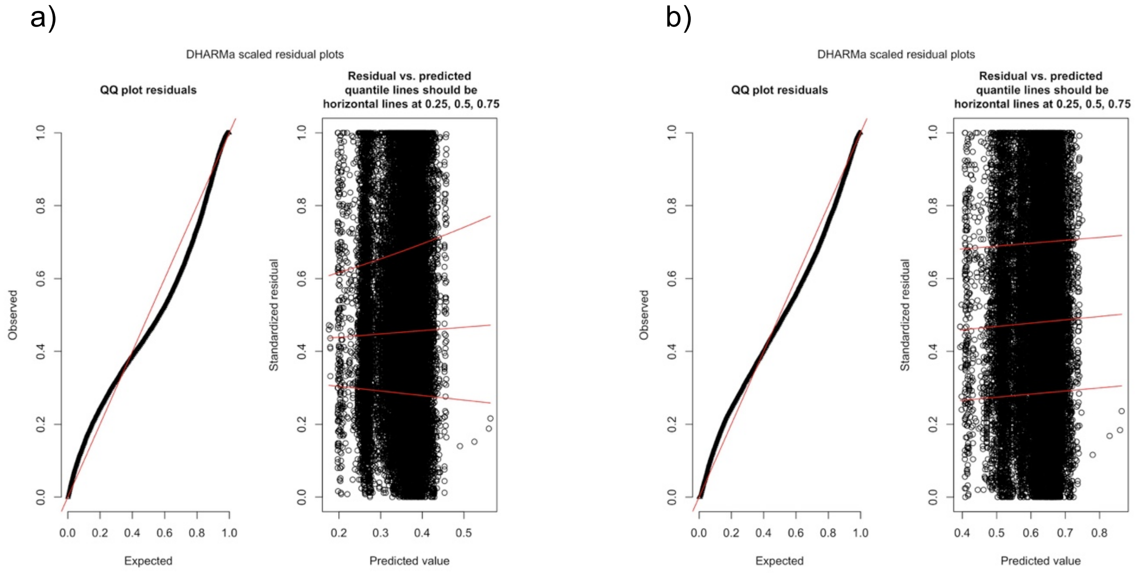


Figure S1. Diagnostic plots for the models using total abundance as response variable. a) MAM using log transformation. b) MAM using square root transformation (final model).

Table S4. AIC values for models with total abundance as response variable using the three different random-effects structures that were tested. Delta AIC values are shown relative to the best model.

| Random-effects structure | d.f. | AIC | delta AIC |
|---|---|---|---|
| (1+LandUse+UseIntensity\|SS)+(1\|SSB) | 122 | -13073.31 | -- |
| (1+LandUse\|SS)+(1\|SSB) | 107 | -12738.73 | 334.58 |
| (1\|SS)+(1\|SSB) | 87 | -10771.61 | 2301.7 |

Table S5. ANOVA table (Type II Wald chisquare tests) for the final model (MAM) using total abundance as response variable

| Term | $\chi^2$ | d.f. | p-value |
|---|---|---|---|
| LUI | 96.23 | 17 | 4.42e-13 *** |
| ISL_MAINL | 0.11 | 1 | 0.74 |
| poly(logHPD, 2) | 25.71 | 2 | 2.61e-06 *** |
| poly(logDistRd, 2) | 15.67 | 2 | 0.0004 *** |
| LUI: ISL_MAINL | 29.28 | 17 | 0.03 * |
| ISL_MAINL: poly(logHPD, 2) | 1.69 | 2 | 0.43 |
| poly(logHPD, 2): LandUse | 47.38 | 10 | 8.06e-07 *** |
| ISL_MAINL: poly(logHPD, 2): LandUse | 48.82 | 10 | 4.40e-07 *** |

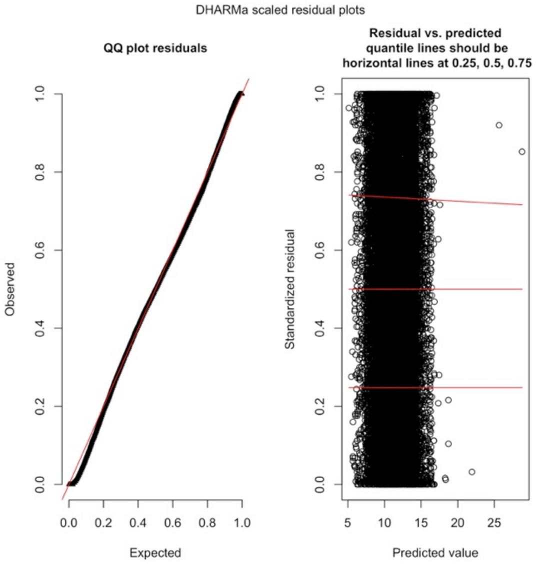


Figure S2. Diagnostic plot for the final model (MAM) using species richness as response variable.

Table S6. AIC values for models with species richness as response variable using the three different random-effects structures that were tested. Delta AIC values are shown relative to the best model.

| Random-effects structure | d.f. | AIC | Delta AIC |
|---|---|---|---|
| (1+LandUse+UseIntensity\|SS)+(1\|SSB) | 121 | 156923.5 | -- |
| (1+LandUse\|SS)+(1\|SSB) | 106 | 157851.2 | 927.7 |
| (1\|SS)+(1\|SSB) | 86 | 163368.6 | 6445.1 |

Table S7. ANOVA table (Type II Wald chisquare tests) for the final model using species richness as response variable.

| Term | $\chi^2$ | d.f. | p-value |
|---|---|---|---|
| LUI | 404.30 | 17 | < 2.2e-16 *** |
| ISL_MAINL | 2.92 | 1 | 0.09 |
| poly(logHPD, 2) | 0.08 | 2 | 0.96 |
| poly(logDistRd, 2) | 15.42 | 2 | 0.0004 *** |
| LUI: ISL_MAINL | 70.29 | 17 | 1.92e-08 *** |
| ISL_MAINL: poly(logHPD, 2) | 28.02 | 2 | 8.24e-07 *** |
| poly(logHPD, 2): LandUse | 147.39 | 10 | < 2.2e-16 *** |
| ISL_MAINL: poly(logDistRd, 2) | 4.31 | 2 | 0.12 |
| LandUse: poly(logDistRd, 2) | 80.11 | 10 | 4.77e-13 *** |
| ISL_MAINL: poly(logHPD, 2): LandUse | 64.25 | 10 | 5.63e-10 *** |
| ISL_MAINL: LandUse: poly(logDistRd, 2) | 55.18 | 10 | 2.92e-08 *** |

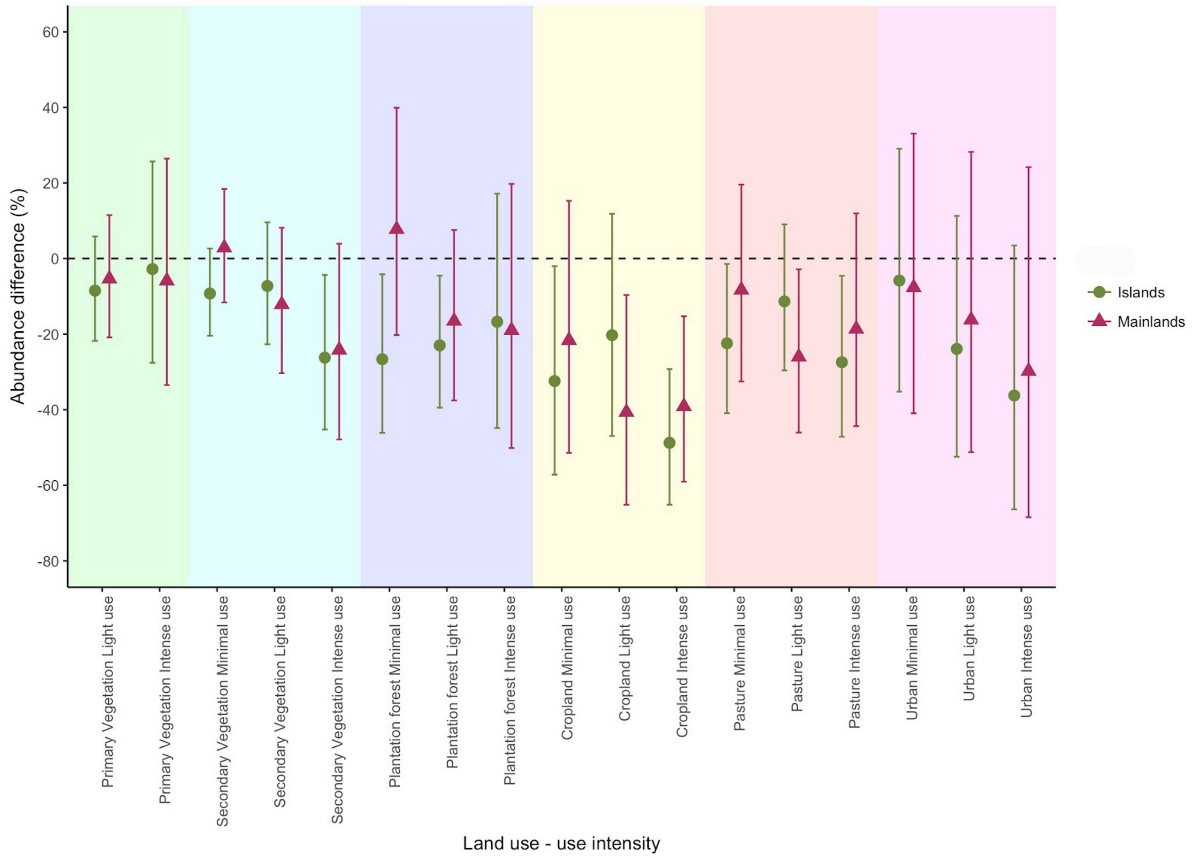


Figure S3. Response of total abundance to LUI on islands and mainlands. Values indicate decrease or increase in percentage of total abundance using minimally-used primary vegetation as baseline. Bars indicate 95% confidence intervals.

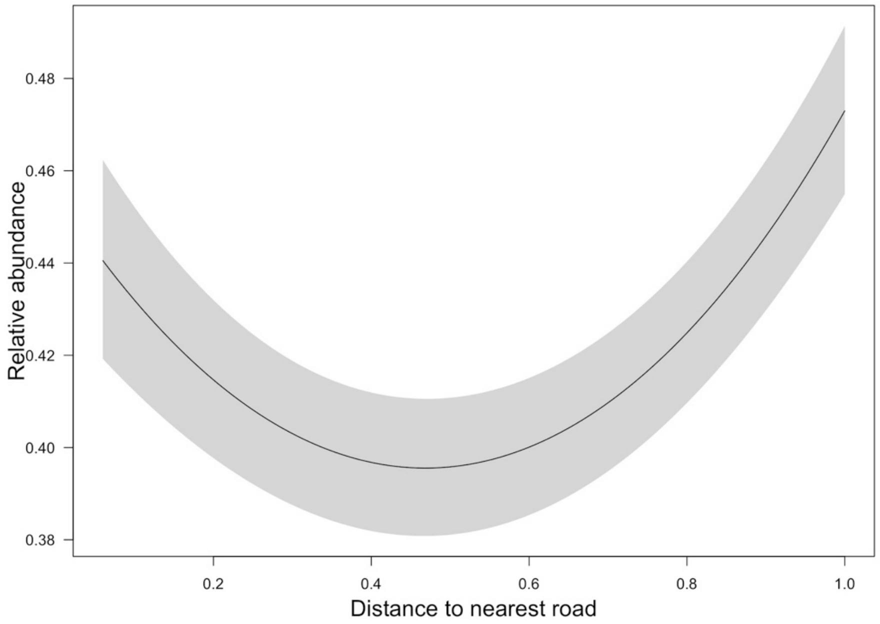


Figure S4. Response of total abundance on island and mainlands to DistRd. DistRd is shown on a rescaled axis (as fitted in the models).

a)

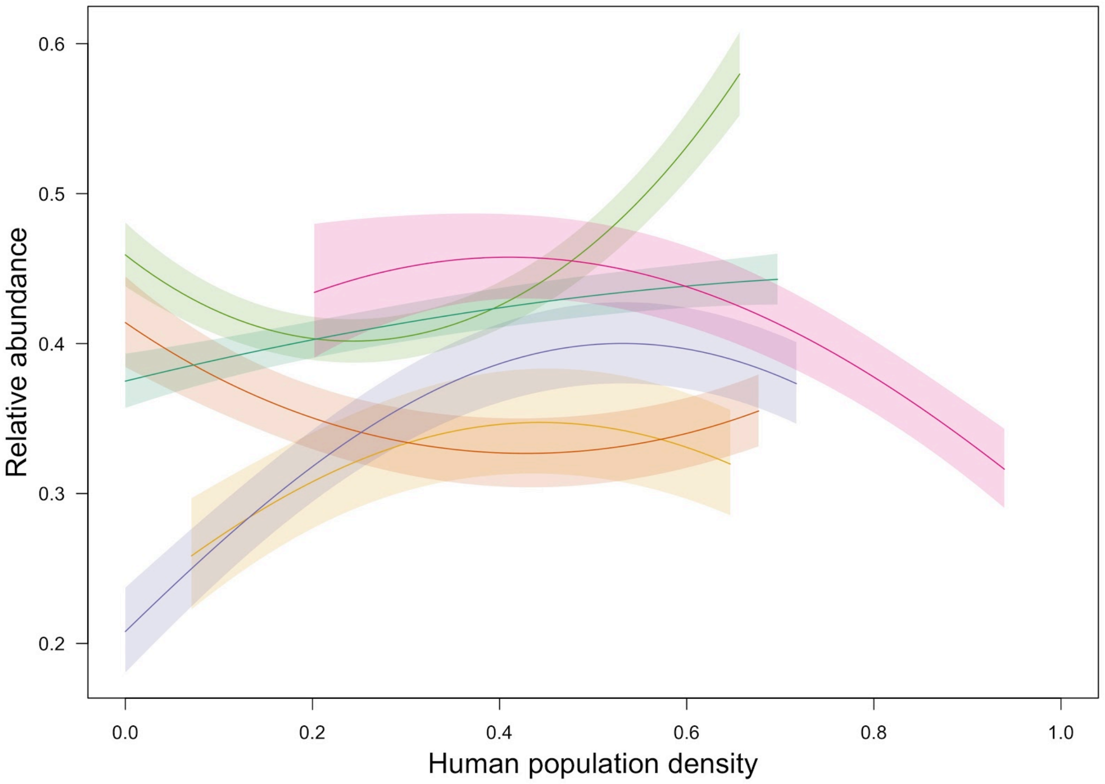


b)

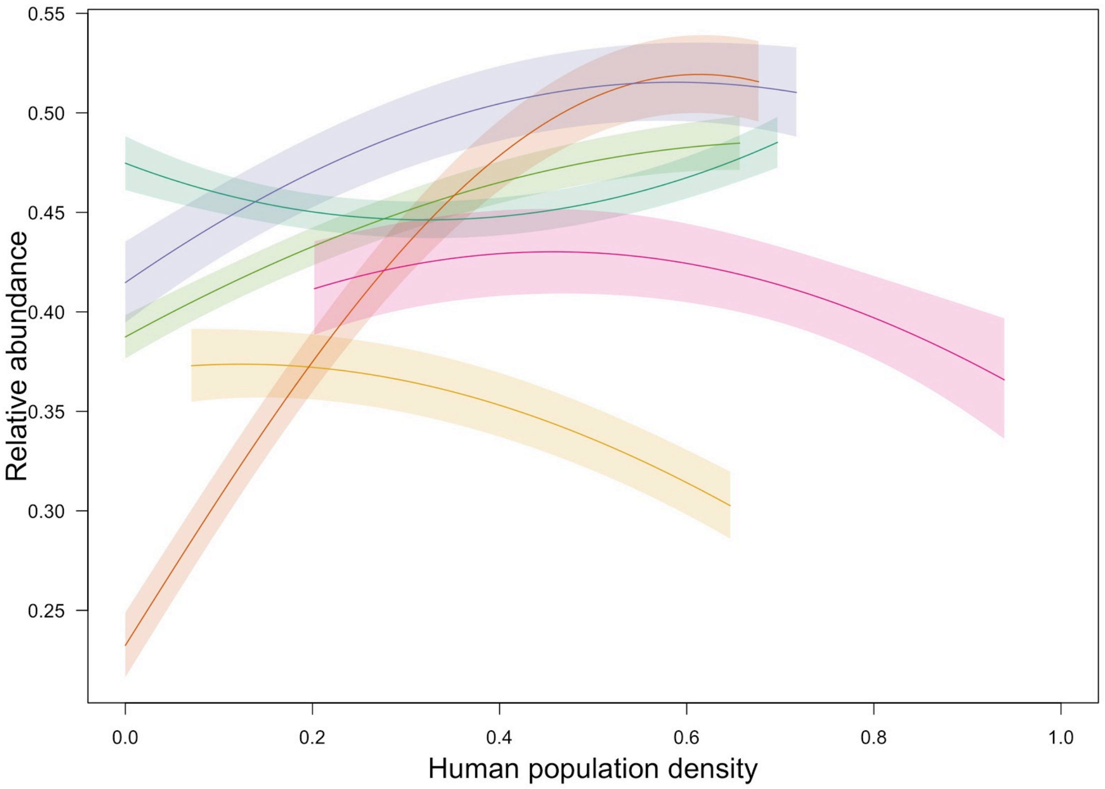


Figure S5. Response of total abundance to HPD in different land uses on a) islands and b) mainlands. HPD is shown on a rescaled axis (as fitted in the models). Land uses are shown in the same colours as in Figure S3.

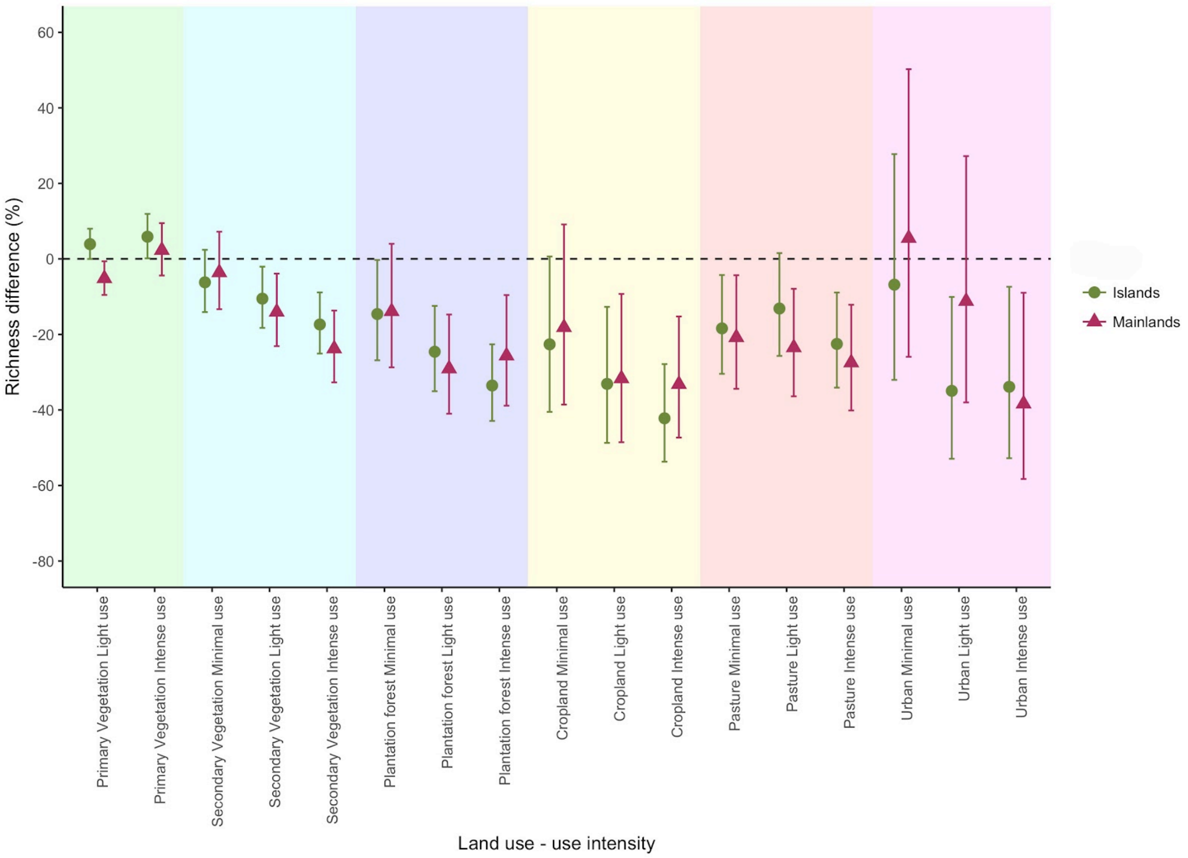


Figure S6. Response of species richness to LUI on islands and mainlands. Values indicate decrease or increase in percentage of species richness using minimally-used primary vegetation as baseline. Bars indicate 95% confidence intervals.

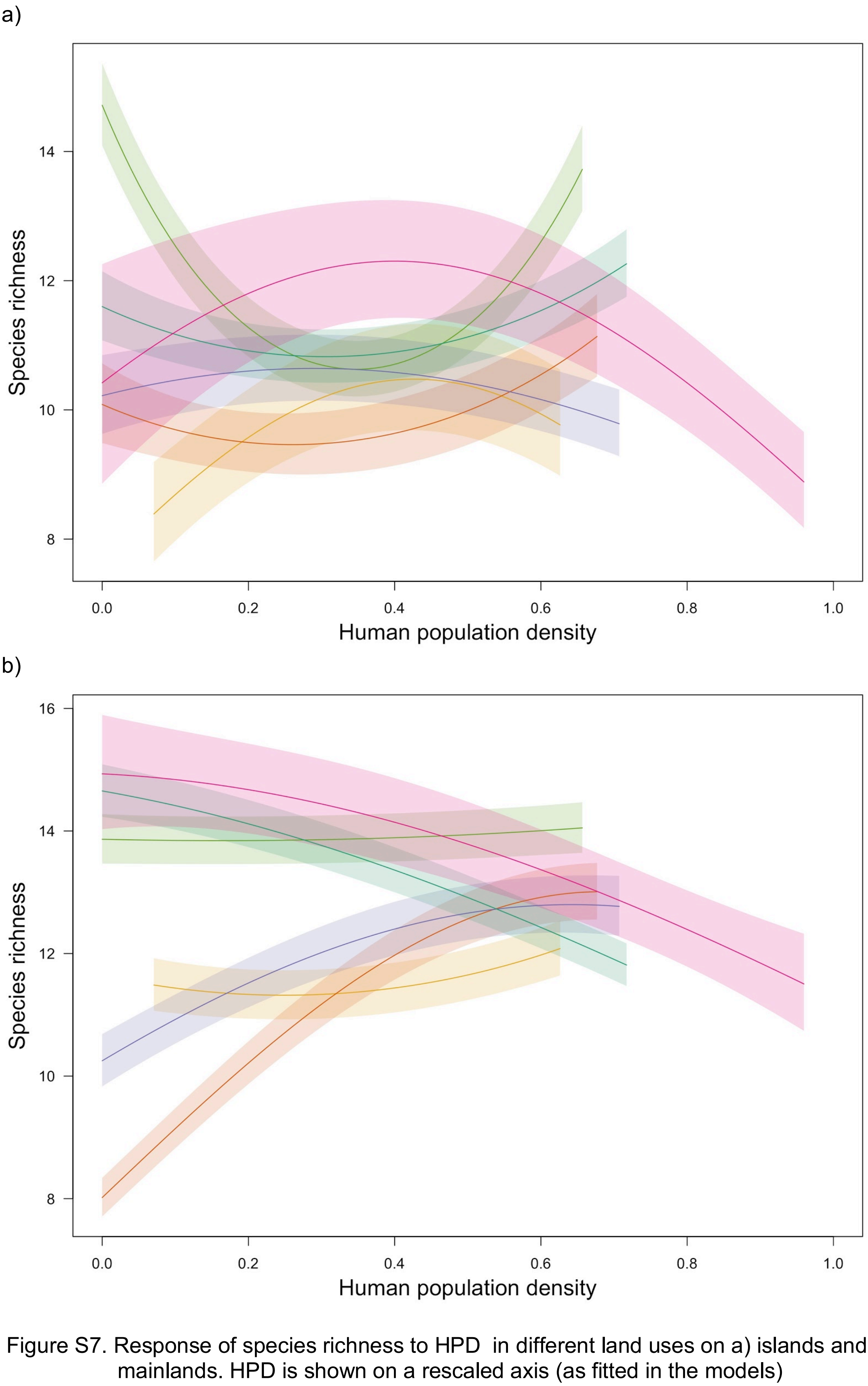


Figure S7. Response of species richness to HPD in different land uses on a) islands and b) mainlands. HPD is shown on a rescaled axis (as fitted in the models)

a)

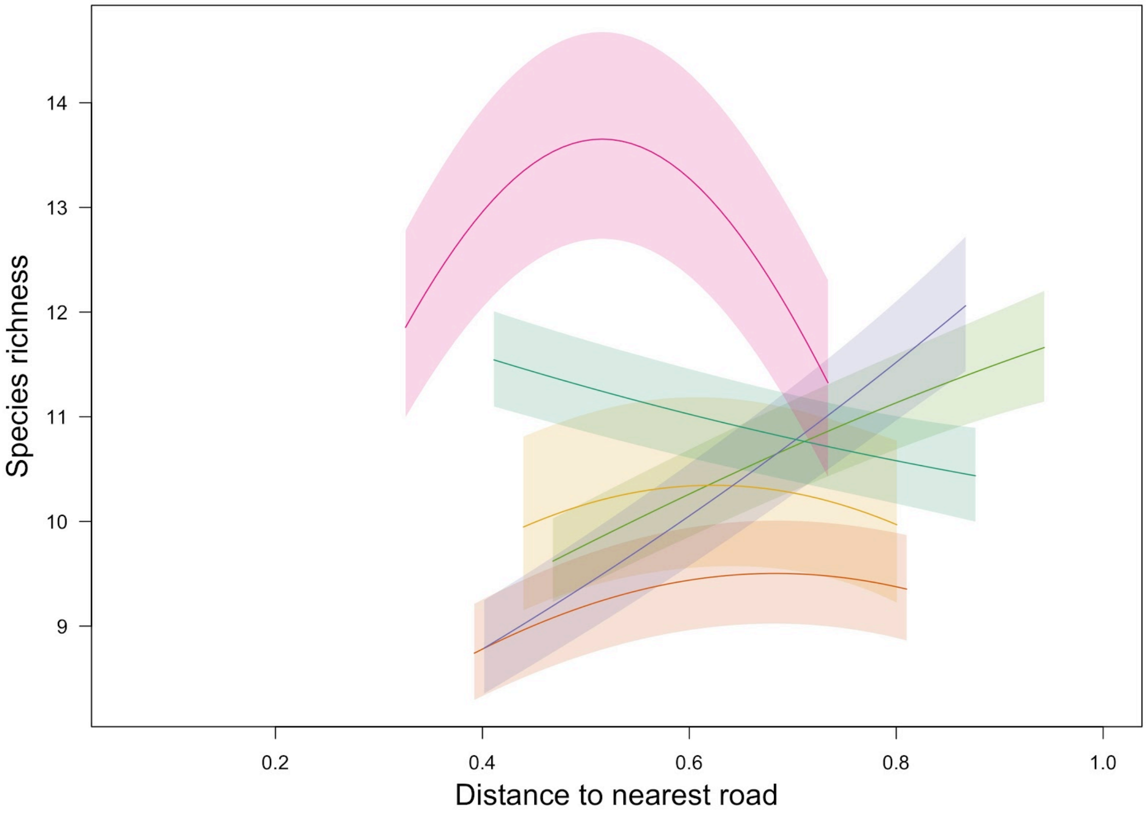


b)

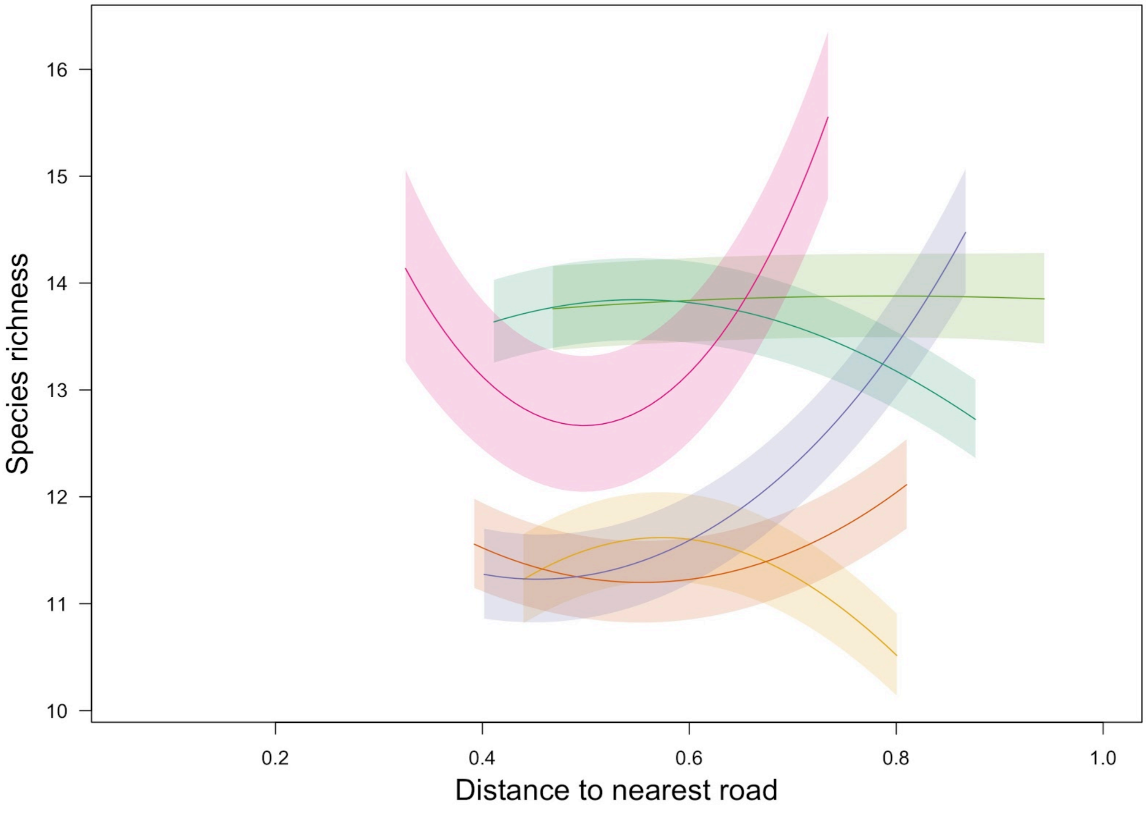


Figure S8. Response of species richness to DistRd in different land uses on a) islands and b) mainlands. DistRd is shown on a rescaled axis (as fitted in the models)

a) b)

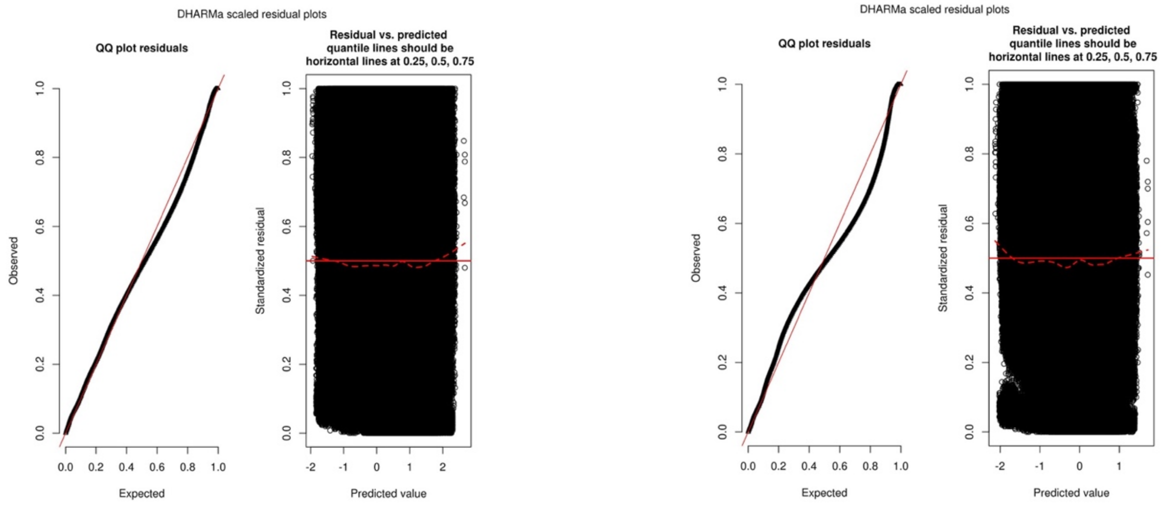


Figure S9. Diagnostic plots for the compositional similarity models. a) Abundance-based and b) Richness-based.

Table S8. Final dataset used for the compositional similarity models. Number of observations per each land-use contrast generated from pairwise comparisons within studies in the PREDICTS database. Numbers in brackets show the number of studies from which data came from. Only land-use contrasts of interest are shown (where site *i* is in PriMin).

| Land-use contrast | Islands | Mainlands |
|---|---|---|
| PriMin- PriMin | 19950 (45) | 391994 (118) |
| PriMin- Primary | 6147 (16) | 15047 (53) |
| PriMin- Secondary | 13757 (38) | 16563 (80) |
| PriMin- Cropland | 4338 (14) | 7375 (23) |
| PriMin- Pasture | 9094 (13) | 19317 (42) |
| PriMin- Plantation | 14494 (27) | 3755 (42) |
| PriMin- Urban | 59 (2) | 8636 (12) |

Table S9. Island and mainland coefficients from the final compositional similarity models (richness and abundance-based). Only results for land-use contrasts with PriMin as baseline are shown. Mainland coefficients are expressed as the difference from the island coefficients. The two coefficients for PriMin-Urb of islands correspond to the original coefficients from the models and coefficients that were calculated indirectly (inside brackets). The coefficients in brackets for PriMin-Urb of mainlands correspond to the difference from the island coefficients that were calculated indirectly. Following the approach to perform backwards stepwise model simplification, we assessed statistical significance of the coefficients corresponding to differences between island and mainland responses (mainland coefficients). We compared the mainland coefficients against null distributions for each coefficient (from 199 null models) to test whether geographic and environmental distance among sites and each land-use contrast had a different effect on compositional similarity when compared to islands (i.e., mainland coefficients that do not differ from the null distributions indicate no real difference from the island coefficients). p-values for each mainland coefficient were calculated by performing a two-tailed hypothesis test. p-values for PriMin-Urban are not shown since they could not be estimated using island coefficients that were calculated indirectly. Significance codes: <0.05**, and 0.005*** (the minimum p-value that can be obtained in our tests is 0.005).

| | Abundance-based model | | Richness-based model | |
|---|---|---|---|---|
| | Islands | Mainlands | Islands | Mainlands |
| PriMin- PriMin | 1.439 | 0.501 *** | 0.603 | 0.403 *** |
| Geographic distance | -0.055 | -0.037 *** | -0.050 | -0.029 *** |
| Environmental distance | -1.511 | 0.534 *** | -1.204 | 0.289 *** |
| PriMin- Primary | -0.460 | -0.079 ** | -0.357 | -0.077 ** |
| PriMin- Secondary | -0.287 | 0.060 | -0.100 | -0.140 *** |
| PriMin- Plantation | -0.745 | -0.740 *** | -0.576 | -0.687 *** |
| PriMin- Cropland | -1.231 | -0.114 ** | -0.920 | -0.142 *** |
| PriMin- Pasture | -1.330 | -0.539 *** | -0.889 | -0.645 *** |
| PriMin- Urban | -0.312 (-1.333) | -1.392 (-0.372) | -0.37 (-1.360) | -0.934 (0.055) |

Table S10. Rescaled backtransformed values (0 to 1 scale) for compositional similarity (richness and abundance-based) of land-use contrasts with PriMin as baseline. Values are expressed relative to compositional similarity between adjacent sites in PriMin with identical environments. For islands, PriMin-Urban compositional similarity inside brackets corresponds to the coefficients that were calculated indirectly.

| | Abundance-based model | | Richness-based model | |
|---|---|---|---|---|
| | Islands | Mainlands | Islands | Mainlands |
| PriMin- PriMin | 1 | 1 | 1 | 1 |
| PriMin- Primary | 0.898 | 0.917 | 0.866 | 0.871 |
| PriMin- Secondary | 0.939 | 0.969 | 0.963 | 0.931 |
| PriMin- Plantation | 0.823 | 0.696 | 0.780 | 0.590 |
| PriMin- Cropland | 0.679 | 0.734 | 0.647 | 0.659 |
| PriMin- Pasture | 0.648 | 0.587 | 0.658 | 0.500 |
| PriMin- Urban | 0.93 (0.647) | 0.635 | 0.86 (0.486) | 0.576 |

Table S11. Results from the compositional similarity model (abundance-based) using log transformation for all variables. The table shows backtransformed values (0 to 1 scale) for compositional similarity of land-use contrasts using PriMin or primary vegetation (primary vegetation with light and intense use) as baseline. Rescaled values are also shown (relative to compositional similarity between adjacent sites in PriMin or primary vegetation with zero environmental distance). We only show results for mainlands as an example.

| | Compositional similarity | Rescaled compositional similarity |
|---|---|---|
| PriMin as baseline | | |
| PriMin-PriMin | 0.674 | 1 |
| PriMin-Primary | 0.633 | 0.939 |
| PriMin-Secondary | 0.659 | 0.978 |
| PriMin-Plantation | 0.585 | 0.868 |
| PriMin-Cropland | 0.586 | 0.870 |
| PriMin-Pasture | 0.566 | 0.840 |
| PriMin-Urban | 0.561 | 0.832 |
| Primary vegetation as baseline | | |
| Primary-Primary | 0.648 | 1 |
| Primary-Secondary | 0.619 | 0.954 |

| | | |
|---|---|---|
| Primary-Plantation | 0.614 | 0.947 |
| Primary-Cropland | 0.590 | 0.910 |
| Primary-Pasture | 0.587 | 0.905 |
| Primary-Urban | 0.617 | 0.951 |

Table S12. Results from the compositional similarity models (abundance-based) when data for PriMin and primary vegetation was collapsed into a single class. The table shows results for two models: using log transformation for all variables and using our alternative transformations, including logit transformation for compositional similarity. Values correspond to backtransformed (0 to 1 scale) compositional similarity values and rescaled values (relative to compositional similarity between adjacent sites in primary vegetation with zero environmental distance). We only show results for mainlands as an example.

| | Logit transformation | | Log transformation | |
|---|---|---|---|---|
| | Compositional similarity | Rescaled compositional similarity | Compositional similarity | Rescaled compositional similarity |
| Primary-Primary | 0.842 | 1 | 0.656 | 1 |
| Primary-Secondary | 0.806 | 0.957 | 0.641 | 0.976 |
| Primary-Plantation | 0.715 | 0.849 | 0.612 | 0.933 |
| Primary-Cropland | 0.642 | 0.762 | 0.587 | 0.893 |
| Primary-Pasture | 0.531 | 0.630 | 0.571 | 0.869 |
| Primary-Urban | 0.635 | 0.754 | 0.588 | 0.895 |

Table S13. Average compositional similarity of all land-use contrasts (using rescaled values) in models using different baselines and data transformations. Only mainland results from the abundance-based models were used to set an example. The cell on the left extreme shows averages from our final abundance-based model (using PriMin as baseline for land-use contrasts and logit transformation for compositional similarity). The cell on the right extreme shows results for the model where the baseline was collapsed primary vegetation (all use intensities) and compositional similarity was log-transformed (as in Newbold et al., 2016). Cells in between show averages for intermediate conditions. The table shows whether the change in baseline or in data transformation caused bigger changes in the range of values for compositional similarity of the different land-use contrasts. We did not include similarity between PriMin-Primary and Primary-PriMin to estimate the averages in order to have the same land-use contrasts across models.

| PriMin + Logit | Primary + Logit | PriMin + Log | Primary + Log |
|---|---|---|---|
| 0.770 | 0.825 | 0.898 | 0.927 |

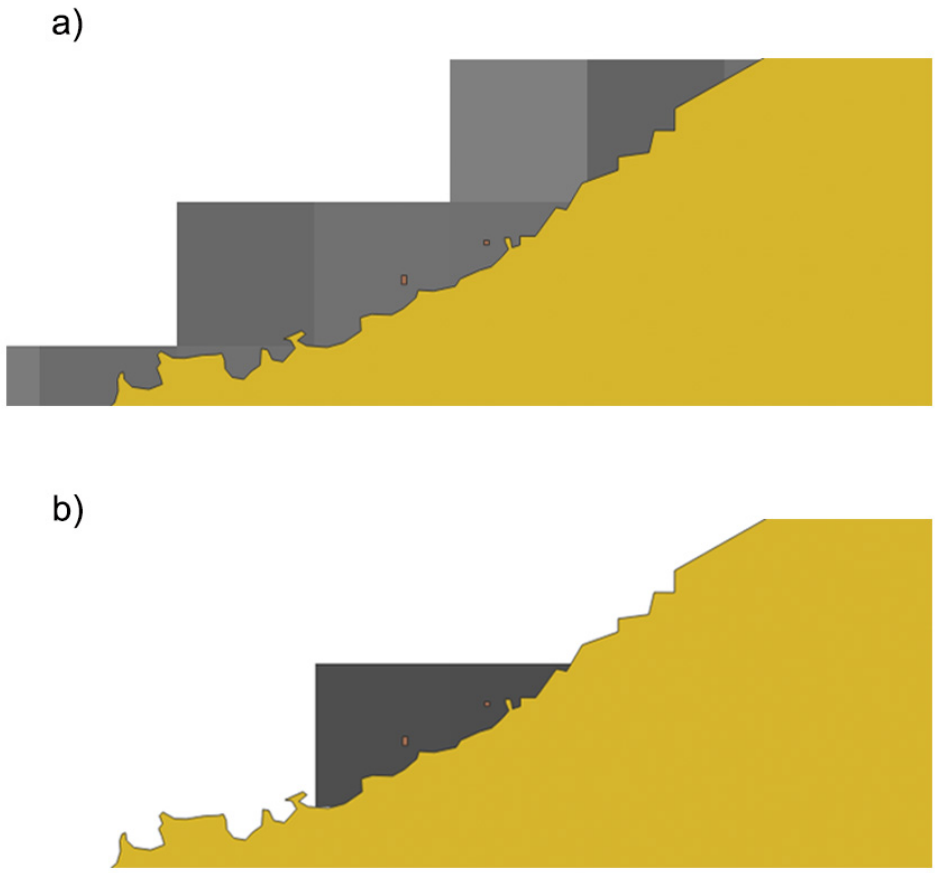


Figure S10. Example of shared cells between islands and mainland in the final BII maps. Two small islands (<1km2) are located close to the mainland coast. The yellow polygons correspond to the mainland shapefile and the small islands are shown in orange. Grey cells show the final BII maps for a) mainlands and b) islands; in this case, two cells of the BII maps are shared between islands and mainlands, since they intersect with both island and mainland shapefiles.

Table S14. Average BII (abundance-based and richness-based) for terrestrial biomes of islands and mainlands. s.d. are shown within parenthesis. BII values are shown in a 0 to 1 scale (1= 100% intactness)

| Biome | Abundance-based BII | | Richness-based BII | |
|---|---|---|---|---|
| | Islands | Mainlands | Islands | Mainlands |
| Temperate Grasslands, Savannas and Shrublands | 0.62 (0.12) | 0.49 (0.13) | 0.52 (0.10) | 0.48 (0.12) |
| Mediterranean Forests, Woodlands and Scrub | 0.59 (0.11) | 0.61 (0.12) | 0.51 (0.09) | 0.57 (0.12) |
| Montane Grasslands and Shrublands | 0.65 (0.08) | 0.60 (0.16) | 0.55 (0.10) | 0.58 (0.15) |
| Tropical and Subtropical Grasslands, Savannas and Shrublands | 0.70 (0.05) | 0.67 (0.13) | 0.59 (0.05) | 0.62 (0.12) |
| Flooded Grasslands and Savannas | 0.63 (0.08) | 0.65 (0.16) | 0.55 (0.10) | 0.61 (0.15) |
| Temperate Broadleaf and Mixed Forests | 0.59 (0.13) | 0.69 (0.13) | 0.52 (0.12) | 0.66 (0.13) |
| Tropical and Subtropical Dry Broadleaf Forests | 0.59 (0.07) | 0.69 (0.11) | 0.49 (0.07) | 0.67 (0.10) |
| Deserts and Xeric Shrublands | 0.70 (0.05) | 0.73 (0.21) | 0.56 (0.04) | 0.71 (0.18) |
| Tropical and Subtropical Coniferous Forests | 0.62 (0.07) | 0.71 (0.12) | 0.52 (0.07) | 0.68 (0.13) |
| Mangroves | 0.71 (0.09) | 0.74 (0.13) | 0.62 (0.11) | 0.70 (0.11) |
| Temperate Conifer Forests | 0.71 (0.11) | 0.76 (0.14) | 0.67 (0.14) | 0.75 (0.15) |
| Tropical and Subtropical Moist Broadleaf Forests | 0.69 (0.09) | 0.83 (0.15) | 0.60 (0.11) | 0.79 (0.15) |
| Boreal Forests/Taiga | 0.77 (0.09) | 0.90 (0.06) | 0.75 (0.11) | 0.92 (0.07) |
| Tundra | 0.96 (0.03) | 0.92 (0.05) | 0.97 (0.04) | 0.97 (0.03) |

Table S15. Average BII (abundance-based and richness-based) for biodiversity hotspots of islands and mainlands. s.d. are shown within parenthesis. BII values are shown in a 0 to 1 scale (1= 100% intactness)

| Hotspot | Abundance-based BII | | Richness-based BII | |
|---|---|---|---|---|
| | Islands | Mainlands | Islands | Mainlands |
| Atlantic Forest | 0.68 (0.06) | 0.67 (0.11) | 0.57 (0.06) | 0.61 (0.11) |
| California Floristic Province | 0.75 (0.1) | 0.69 (0.17) | 0.65 (0.12) | 0.68 (0.18) |
| Cape Floristic Region | -- | 0.45 (0.12) | -- | 0.44 (0.1) |
| Caribbean Islands | 0.58 (0.06) | -- | 0.49 (0.07) | -- |
| Caucasus | 0.69 (0.06) | 0.64 (0.12) | 0.57 (0.04) | 0.6 (0.11) |
| Cerrado | -- | 0.62 (0.12) | -- | 0.59 (0.11) |
| Chilean Winter Rainfall and Valdivian Forests | 0.77 (0.05) | 0.78 (0.12) | 0.65 (0.05) | 0.76 (0.12) |
| Coastal Forests of Eastern Africa | 0.56 (0.08) | 0.72 (0.09) | 0.49 (0.05) | 0.65 (0.08) |
| East Melanesian Islands | 0.74 (0.04) | -- | 0.66 (0.3) | -- |
| Eastern Afromontane | -- | 0.75 (0.1) | -- | 0.68 (0.1) |
| Guinean Forests of West Africa | 0.66 (0.04) | 0.74 (0.1) | 0.57 (0.03) | 0.68 (0.1) |
| Himalaya | -- | 0.68 (0.16) | -- | 0.65 (0.14) |
| Horn of Africa | 0.6 (0.07) | 0.66 (0.1) | 0.49 (0.06) | 0.61 (0.1) |
| Indo-Burma | 0.64 (0.09) | 0.79 (0.13) | 0.52 (0.08) | 0.77 (0.12) |
| Irano-Anatolian | -- | 0.66 (0.09) | -- | 0.6 (0.09) |
| Japan | 0.64 (0.09) | -- | 0.55 (0.08) | -- |
| Madagascar and the Indian Ocean Islands | 0.58 (0.06) | -- | 0.49 (0.05) | -- |
| Madrean Pine-Oak Woodlands | -- | 0.7 (0.13) | -- | 0.67 (0.14) |
| Maputaland-Pondoland-Albany | 0.63 (0.06) | 0.59 (0.15) | 0.5 (0.06) | 0.56 (0.12) |
| Mediterranean Basin | 0.55 (0.08) | 0.61 (0.11) | 0.47 (0.07) | 0.57 (0.11) |
| Mesoamerica | 0.68 (0.06) | 0.75 (0.1) | 0.57 (0.06) | 0.7 (0.12) |
| Mountains of Central Asia | -- | 0.61 (0.12) | -- | 0.56 (0.12) |
| Mountains of Southwest China | -- | 0.66 (0.13) | -- | 0.61 (0.13) |
| New Caledonia | 0.7 (0.04) | -- | 0.68 (0.07) | -- |
| New Zealand | 0.58 (0.1) | -- | 0.5 (0.1) | -- |
| Philippines | 0.62 (0.07) | -- | 0.5 (0.06) | -- |

| | | | | |
|---|---|---|---|---|
| Polynesia-Micronesia | 0.64 (0.09) | -- | 0.54 (0.1) | -- |
| Southwest Australia | 0.6 (0.1) | -- | 0.52 (0.07) | -- |
| Succulent Karoo | 0.67 (0.03) | 0.4 (0.14) | 0.57 (0.05) | 0.43 (0.13) |
| Sundaland | 0.68 (0.09) | 0.8 (0.11) | 0.59 (0.11) | 0.78 (0.1) |
| Tropical Andes | 0.72 (0.01) | 0.77 (0.1) | 0.58 (0.01) | 0.74 (0.12) |
| Tumbes-Choco-Magdalena | 0.8 (0.09) | 0.76 (0.1) | 0.75 (0.12) | 0.71 (0.1) |
| Wallacea | 0.69 (0.06) | -- | 0.59 (0.08) | -- |
| Western Ghats and Sri Lanka | 0.66 (0.07) | 0.75 (0.13) | 0.54 (0.06) | 0.72 (0.09) |

Table S16. Percentage of land surface with each LUI in the final BII island (including and excluding Australia) and mainland maps.

| LUI | Percentage on islands | Percentage on mainlands | Percentage on islands excluding Australia |
|---|---|---|---|
| Primary Vegetation Minimal use | 38.45 | 28.48 | 45.21 |
| Primary Vegetation Light use | 8.35 | 8.79 | 14.40 |
| Primary Vegetation Intense use | 1.03 | 0.94 | 2.07 |
| Secondary Vegetation Minimal use | 7.40 | 16.75 | 8.21 |
| Secondary Vegetation Light use | 2.29 | 6.15 | 3.84 |
| Secondary Vegetation Intense use | 0.61 | 1.52 | 1.22 |
| Cropland Minimal use | 1.14 | 2.65 | 1.91 |
| Cropland Light use | 2.99 | 4.24 | 4.19 |
| Cropland Intense use | 4.85 | 5.40 | 5.67 |
| Pasture Minimal use | 0.29 | 0.11 | 0.25 |
| Pasture Light use | 29.74 | 21.61 | 9.62 |
| Pasture Intense use | 2.35 | 2.96 | 2.59 |
| Urban Minimal use | 0.32 | 0.23 | 0.54 |
| Urban Light use | 0.01 | 0.0007 | 0.01 |
| Urban Intense use | 0.16 | 0.16 | 0.29 |

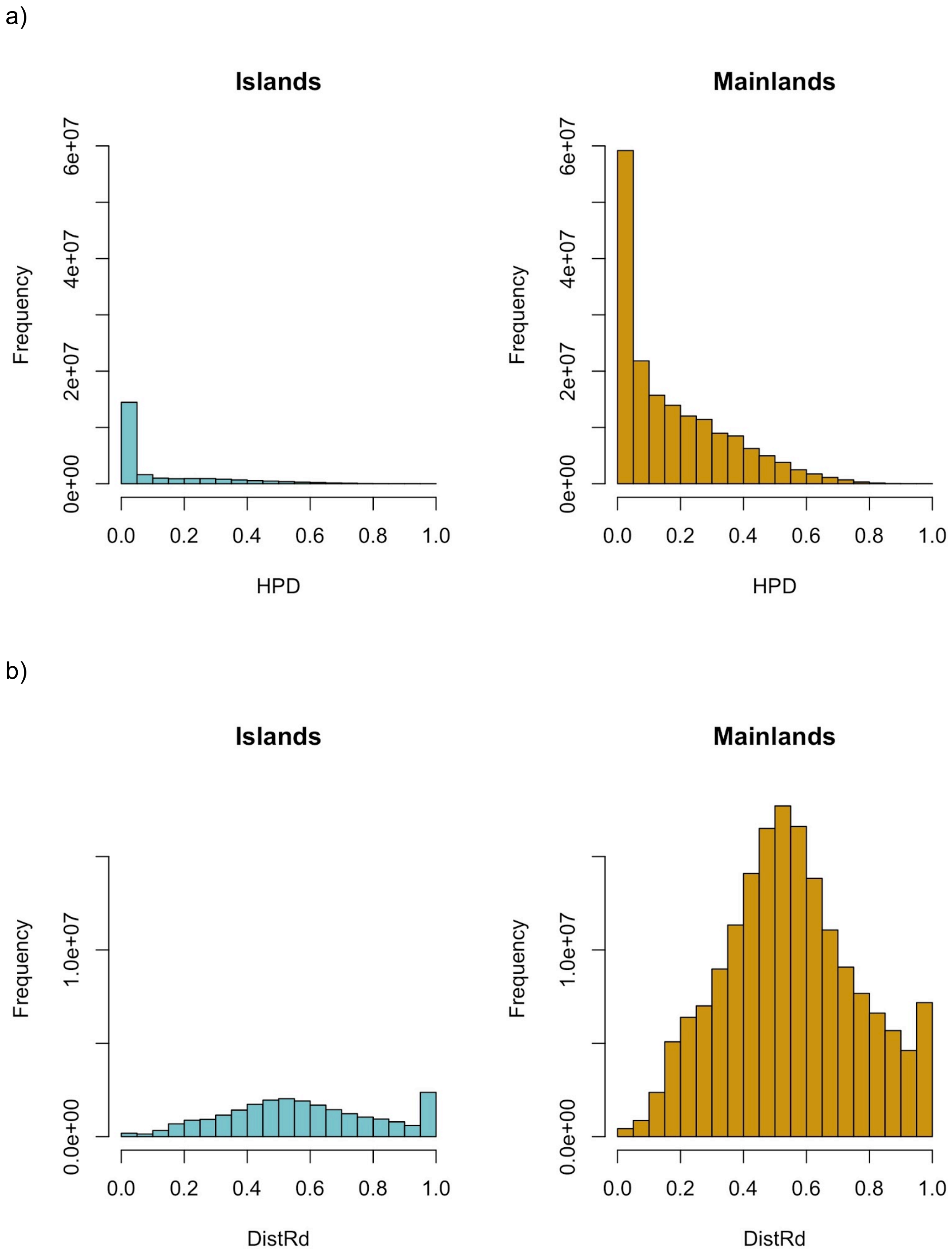


Figure S11. Data distribution for a) human population density (HPD) and b) distance to the nearest road (DistRd) on island and mainland maps. Frequency indicates number of cells in the maps. Only cells that had a defined value in the final BII maps are included. HPD and DistRd are shown on rescaled axes (as fitted in the models). The high frequency of cells with a high value for distance to road might evidence the lack of data in gROADS maps for some regions, especially for islands.

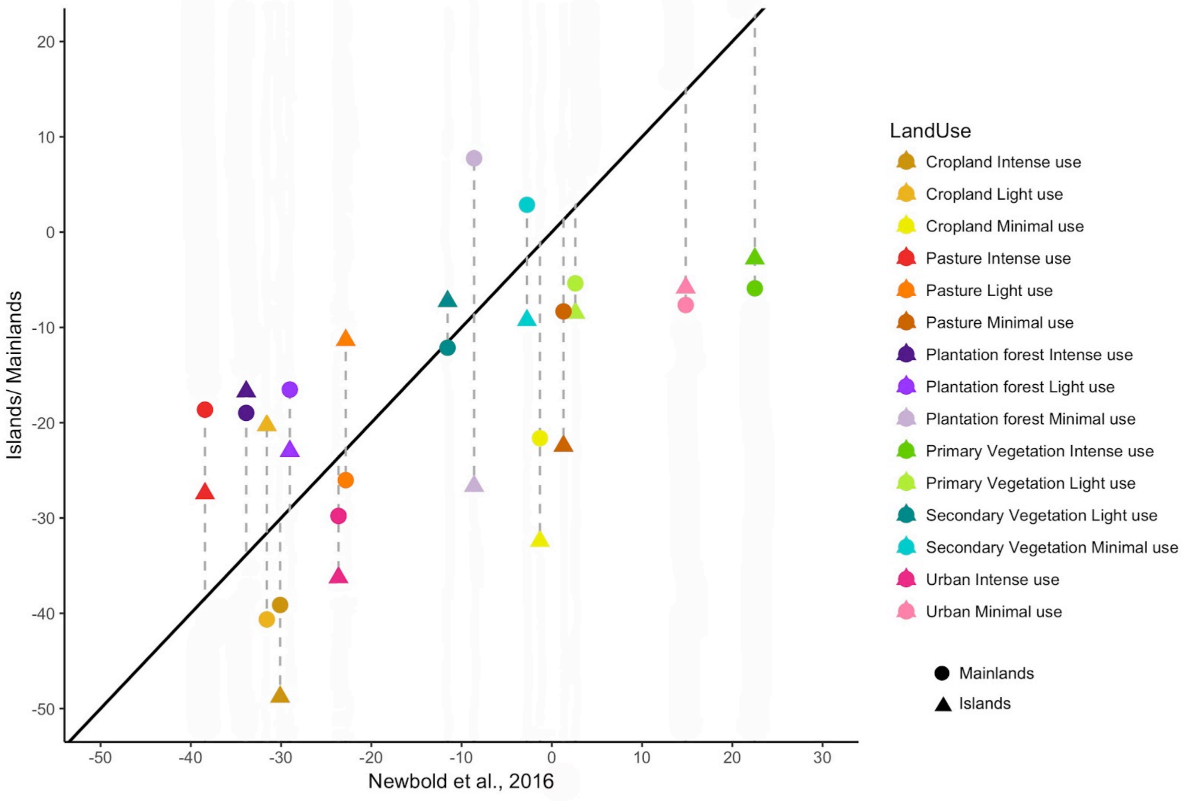


Figure S12. Responses of total abundance to LUI. Island and mainland estimates are compared against global estimates from Newbold et al. (2016). Values indicate decrease or increase in percentage of total abundance using minimally-used primary vegetation as baseline.

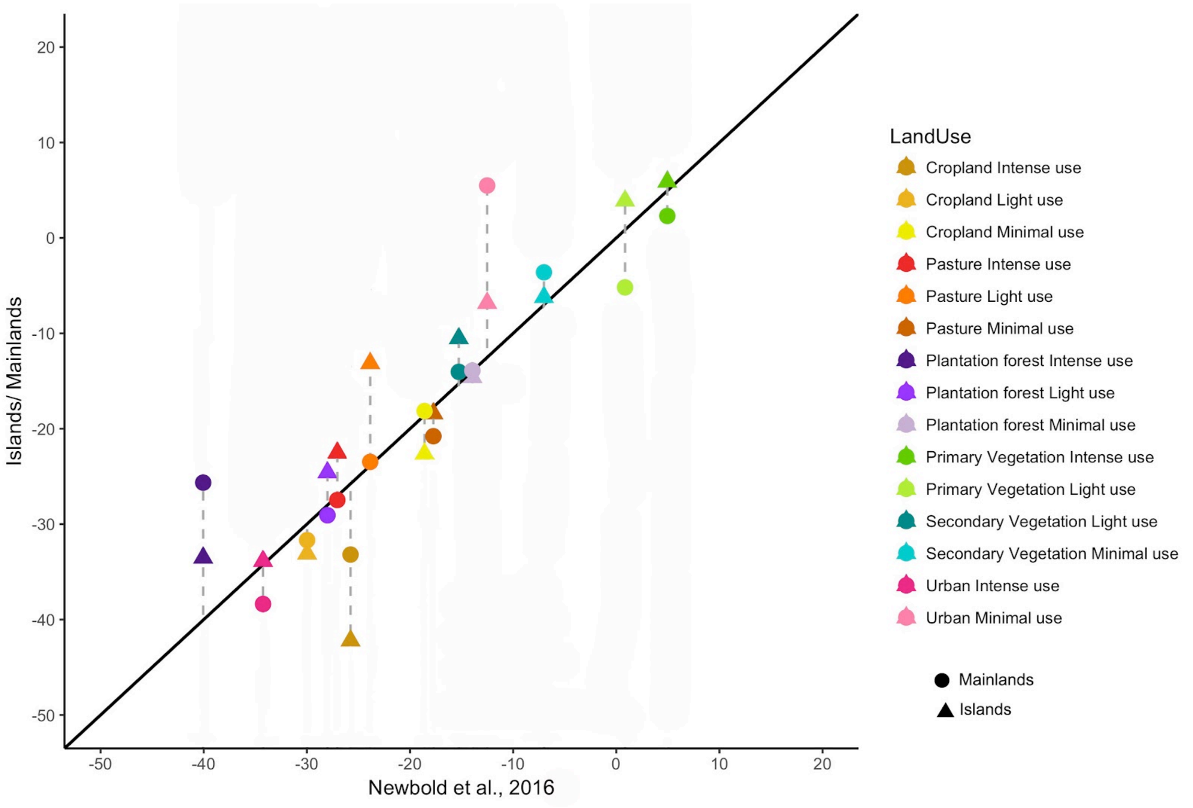


Figure S13. Responses of species richness to LUI. Island and mainland estimates are compared against global estimates from Newbold et al. (2016). Values indicate decrease or increase in percentage of species richness using minimally-used primary vegetation as baseline.

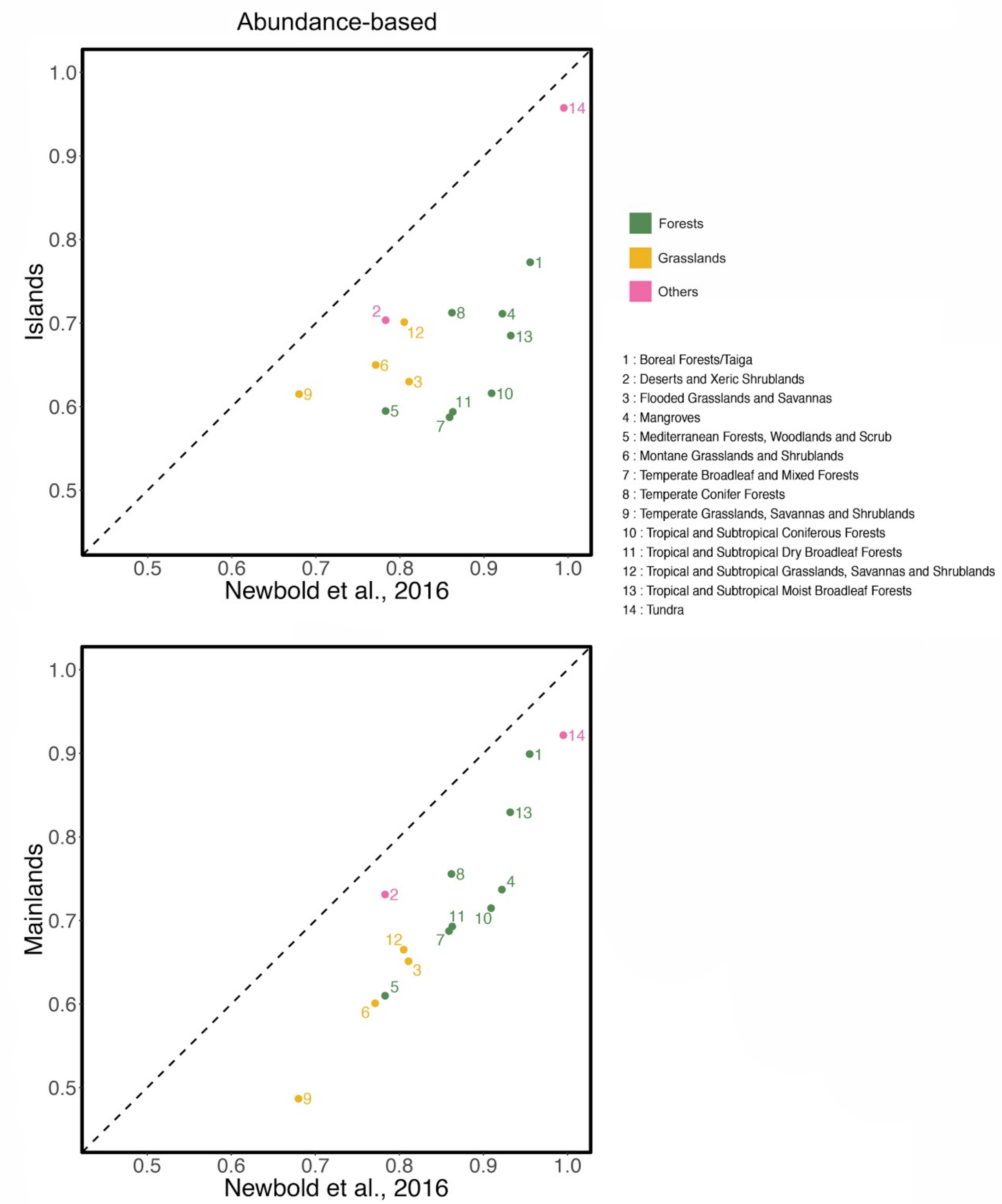


Figure S14. Average biodiversity intactness for the different biomes (abundance-based). Island and mainland averages are compared to averages from Newbold et al. (2016). Colours indicate major biome type. Values from 0 to 1 correspond to BII (1= 100% intactness)

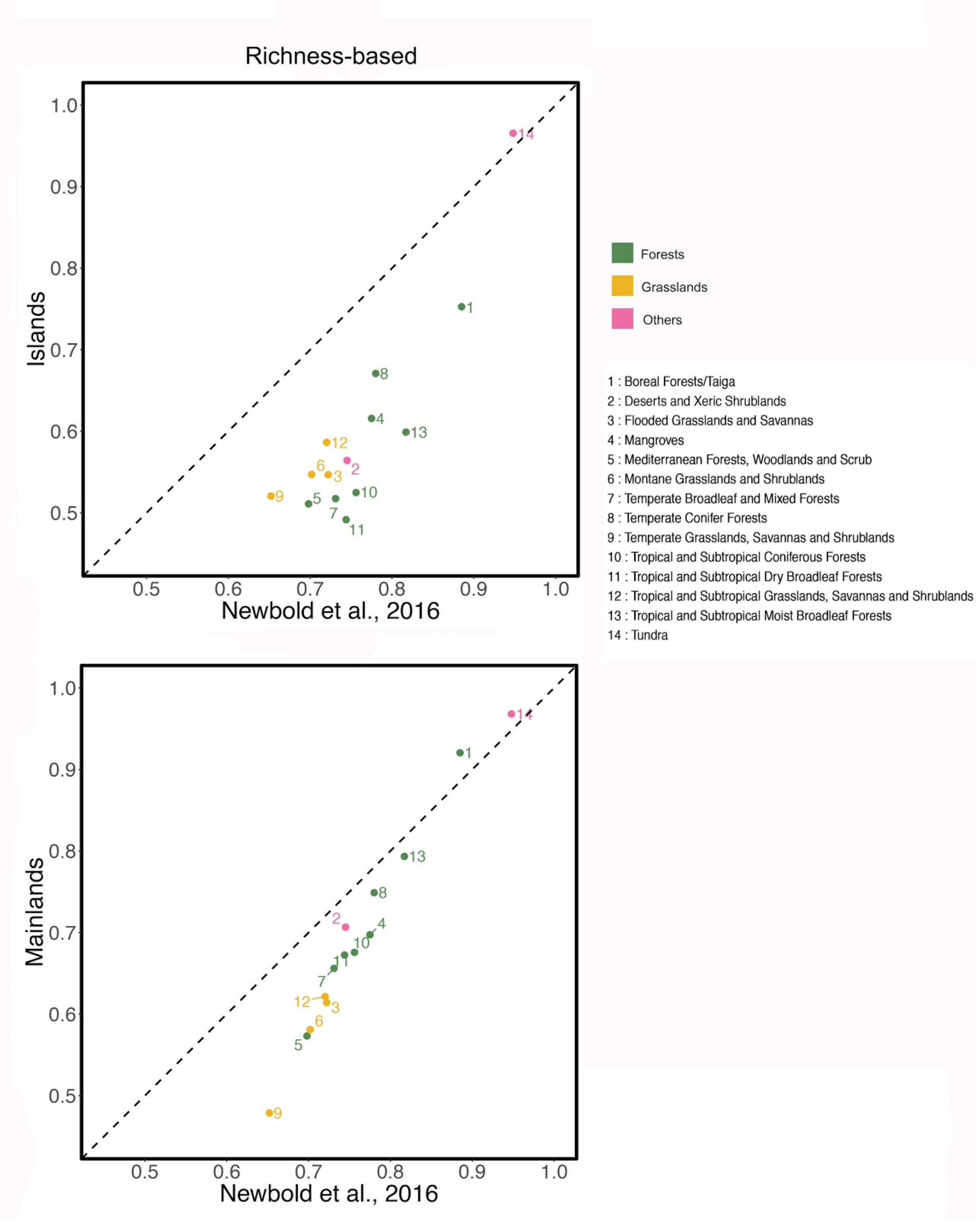


Figure S15. Average biodiversity intactness for different biomes (richness-based). Island and mainland averages are compared to averages from Newbold et al. (2016). Colours indicate major biome type. Values from 0 to 1 correspond to BII (1= 100% intactness)

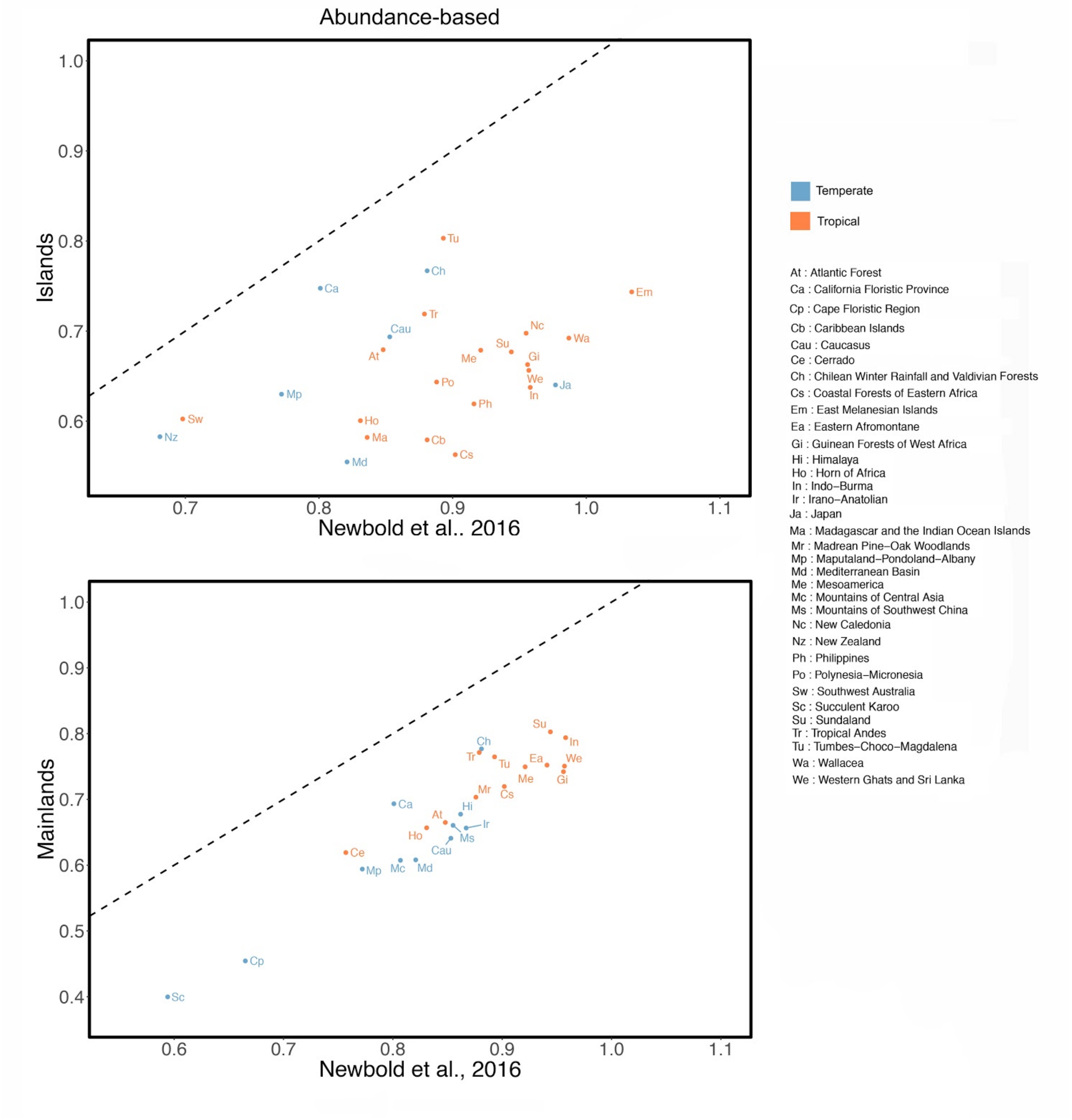


Figure S16. Average biodiversity intactness for the different biodiversity hotspots (abundance-based). Island and mainland averages are compared to global averages from Newbold et al. (2016). Some hotspots are exclusively located on islands or mainlands. Colours indicate whether hotspots are in the tropical or temperate realms. Values from 0 to 1 correspond to BII (1= 100% intactness)

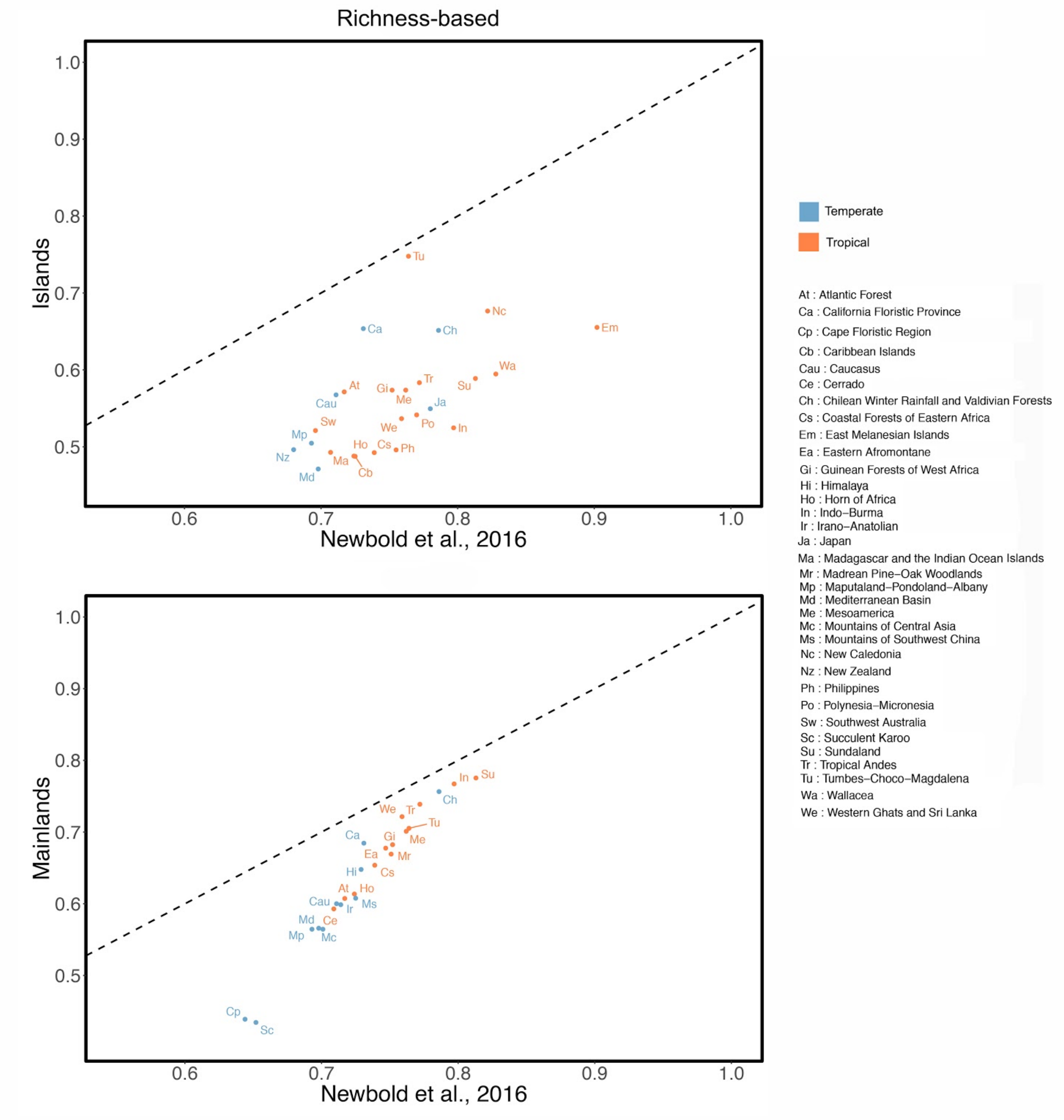


Figure S17. Average biodiversity intactness for the different biodiversity hotspots (richness-based). Island and mainland averages are compared to global averages from Newbold et al. (2016). Some hotspots are exclusively located on islands or mainlands. Colors indicate whether hotspots are in the tropical or temperate realms. Values from 0 to 1 correspond to BII (1= 100% intactness)

a)

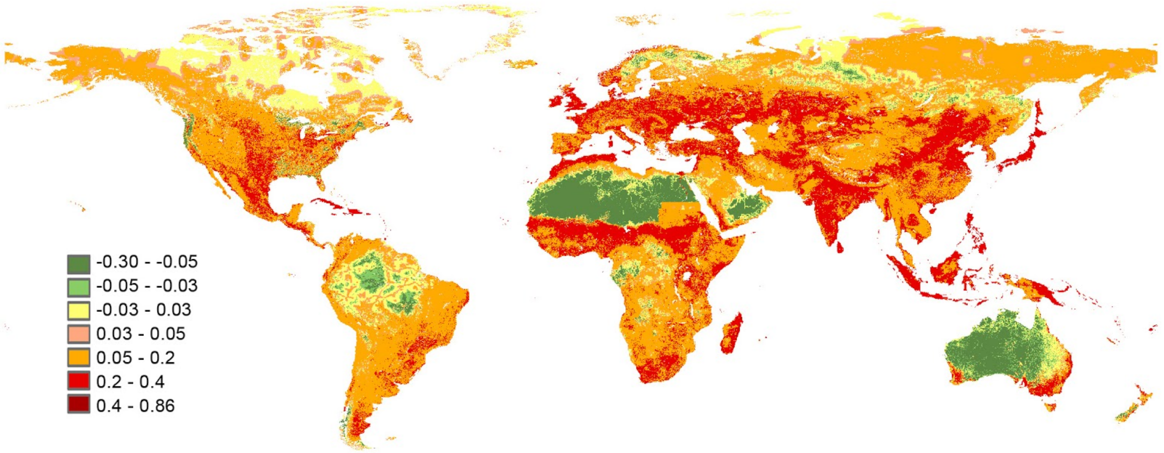


b)

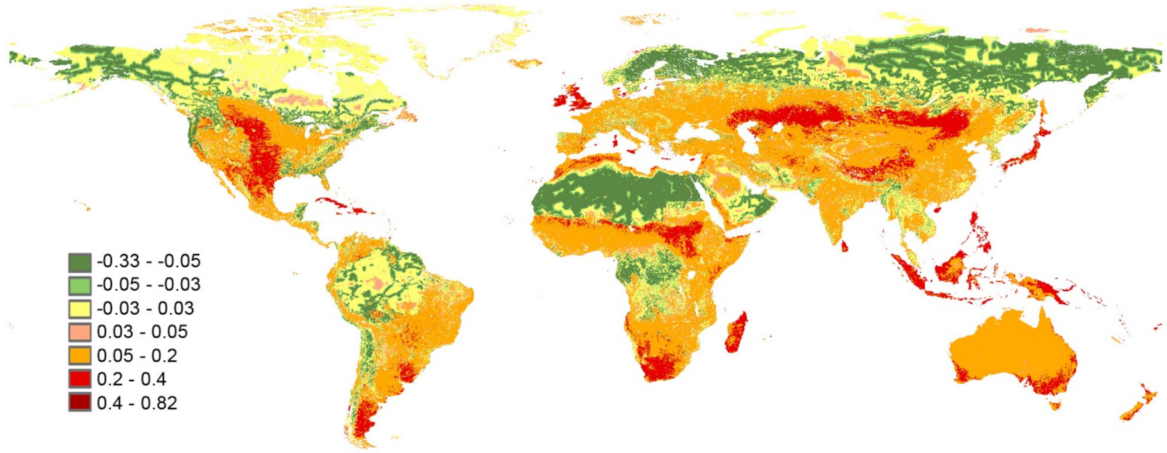


Figure S18. Differences between BII values from Newbold et al. (2016) and the current study. Maps were produced by subtracting our global BII raster (islands + mainlands) from the global raster from Newbold et al. Positive values (shown in red and orange) indicate cases where our estimates are lower than those in Newbold et al. Negative values (shown in green) indicate cases where our estimates are higher than previously. The yellow areas show cases where estimates from both studies differ minimally (maximum by 0.03). a) Abundance-based, b) Richness-based.
